# Organoid modeling of tumor-associated macrophages reveals phagocytosis checkpoint blockade-induced conversion to an immunosuppressive SPP1+ phenotype

**DOI:** 10.64898/2026.05.06.722767

**Authors:** Michitaka Nakano, Lyong Heo, Yu-Ping Yang, Lorena P. Munoz, Yihua Liu, Lei Zhao, Juhyung Park, Eirini Tsekitsidou, Anthony Francois, Jie Liu, Aaron C. Trotman-Grant, Maria Fernanda Henao Echeverri, Cara C. Rada, Edward Tran, Arya Khokhar, Kanako Yuki, Asmita Bhattacharya, Hudson T. Horn, Roel Polak, Leonard N. Yenwongfai, Ying Li, Matt Peach, Emon Nasajpour, Ana Jimena Pavlovitch-Bedzyk, Anne Lynn S. Chang, Michael Lim, Claudia K. Petritsch, Melanie Hayden Gephart, John T. Leppert, Ramesh V. Nair, Mark M. Davis, Michael C. Bassik, Melody Zhang, Jared Odegard, Jamie G. Bates, Lawrence L. K. Leung, Ravindra Majeti, Calvin J. Kuo

## Abstract

Tumor-associated macrophages (TAM) exert essential functions during the immune response to cancer. However, investigations of TAM within a native human tumor microenvironment (TME) have been impeded by a lack of appropriate model systems. Here, patient-derived organoids (PDO) from air-liquid interface (ALI)-grown tumor fragments, containing a human TME that encompassed stroma and immune subsets, robustly preserved TAM that were maintained by endogenous CSF-1 and appropriately responded to polarization signals. Antibody blockade of the CD47 regulatory checkpoint in organoids stimulated phagocytosis and remodeled TAM cytokine secretion profiles that were confirmed in anti-CD47 phase I trial patients. Amongst PDO histologies screened, anti-CD47 tumor killing was notable in clear cell renal cell carcinoma (ccRCC) which was associated with increased TAM infiltration. PDO contained diverse previously described TAM subsets; however, anti-CD47 reprogrammed organoid TAM toward an immunosuppressive SPP1+ phenotype, highlighting a negative feedback mechanism. Our findings uncover a resistance circuit engaged by macrophage checkpoint blockade and position ALI PDO as a robust translational platform for dissecting human macrophage biology and informing precision immunotherapy.

## Introduction

The tumor microenvironment (TME) encompasses stroma and diverse immune subsets, whose manipulation holds substantial promise for cancer treatment (1–3).Within the immune TME, tumor-associated macrophages (TAM) have emerged as a target for cancer immunotherapy (4–6). TAM exert central functions within the TME, residing at the nexus of cancer cell phagocytosis, antigen presentation and immune network interactions(7–9). However, contemporary *in vitro* tumor culture systems lack TAM and thus do not recapitulate these complex TAM functions, hindering mechanistic investigations and development of TAM-directed cancer therapies.

TAMs have been historically characterized as M1 pro-inflammatory TAMs that exert anti-tumor effects, or M2 anti-inflammatory and pro-tumorigenic TAMs (10,11). However, recent single cell multiomic studies have revealed additional TAM phenotypic and functional diversity (12–16). *SPP1+* TAM support fibrosis and matrix remodeling (13,17,18), are associated with poor prognosis in human cancer (19,20), and have attracted significant interest because of their association with immunosuppression (21–24) and cancer immunotherapy resistance (25,26). *C1Q+* TAMs express complement components and MHC class II molecules (27,28), promote tumor progression(28,29) and are mutually exclusive with *SPP1+* TAM (30,31). The *NLRP3+* TAM subset is defined by inflammasome expression (32), releasing pro-inflammatory cytokines such as IL-1β in response to innate stimuli (33). *CXCL9+* TAM are regulated by IFN-γ signaling and correlate with immune checkpoint inhibition response. Together, these individual TAM subsets embody distinct functions within the TME(31,34).

Tumor phagocytosis is a canonical TAM activity with potential for cancer therapeutic manipulation. Tumor cells can escape phagocytosis by elaborating cell surface “don’t eat me” signals such as CD47 (35), CD24 (36), and MHC-I, which interact with negative regulatory checkpoint inhibitory molecules on TAM, such as signal protein-α (SIRPα), sialic acid-binding immunoglobulin-like lectin (SIGLEC-10) and leucocyte immunoglobulin-like receptor B1 (LILRB1), respectively. As CD47 is ubiquitously expressed in normal cells and overexpressed in diverse cancers (37–39), promoting TAM-mediated tumor phagocytosis by antibody blockade of CD47-SIRPα has been investigated for cancer therapy (40–42). However, clinical trials of blocking CD47 or SIRPα with or without additional agents have not demonstrated efficacy or safety in hematologic malignancies (43–46), while solid tumor activity (47–49) remains to be confirmed in definitive trials. To realize the potential of TAM-targeted therapies in general, further explorations are needed to define biological mechanisms, optimize response, resistance and safety, and stratify responsive patients by prognostic biomarkers and tumor histologies.

Patient-derived tumor organoids (PDO) are widely used for three-dimensional human tumor culture, but typically only contain tumor epithelium and lack stromal and immune components (50–53). Recent advances in tumor immunotherapy have incurred a pressing need to extend PDO technology beyond epithelial tumor compartments to include immune subsets that would model the complex immune-tumor intratumoral crosstalk (54–57). We previously reported a 3D air-liquid interface (ALI) organoid method that cultures intact human tumor fragments with endogenous immune components (58,59). Here, we address the longstanding need for *in vitro* modeling of TME macrophages by demonstrating that ALI tumor organoids robustly preserve TAM subsets that are functionally-responsive and phagocytosis-competent, and exploit this system for mechanistic assessment of TAM-targeted anti-tumor therapies, using CD47 inhibition as proof-of-principle.

## Results

### Human tumor ALI organoids preserve TAM within a holistic immune TME

We used an ALI system to culture intact fragments of surgically resected human cancer tissues. The PDO were embedded in a collagen matrix within a transwell exposed directly to air and cultured in media containing WNT3A, EGF, Noggin, R-spondin-1 (WENR) and low-dose IL-2 in an outer dish to support tumor and immune cells **(see Methods)**. Across all experiments, organoids were created from surgical tumor fragments from 67 clear cell renal cell carcinoma (ccRCC), 46 non-small cell lung cancer (NSCLC), and 31 colorectal adenocarcinoma (CRC) patients, but also including additional tumor histologies **(Fig. 1A, Supplementary Table S1)**. PDO cultures typically reproduced tumor architecture with distinct epithelial and stromal compartments (**Fig. 1B-D**). Immunofluorescence (IF) demonstrated IBA1+ TAM across all ALI organoid cancer types surveyed, in association with tumor epithelium appropriately expressing CA9 (ccRCC) (**Fig. 1E**), CK7 (NSCLC adenocarcinoma) (**Fig. 1F**) or CK19 (CRC) (**Fig. 1G**). Upon flow cytometry, CD163 and scatter characteristics were used to differentiate TAM versus smaller monocytes lacking CD163 (62). ALI organoids preserved CD45+CD11b+HLA-DR+CD14+CD163+ TAM compared to freshly isolated tumor specimens at culture days 7, 14 and 28 (**Fig. 1H, Supplementary Fig. S1A**). The detected degree of TAM preservation could be influenced by concomitant tumor cell proliferation. In contrast, PDO CD45+CD11b+HLA-DR+CD14+CD163-monocytes rapidly decreased, consistent with their short in vivo lifespan of 1-7 days (63) (**Supplementary Fig. S1A-B**). Organoid TAM were still detected by histology at extended time points (85 days was the longest time attempted), albeit at significantly decreased and/or variable levels (**Supplementary Fig. S1C)**. ALI PDO could be cryopreserved in-gel and recovered, preserving tissue architectures and immune subsets including TAM (**Supplementary Fig. S1D-F).** We also evaluated PDO preservation of tumor identity by targeted DNA sequencing of selected cases to confirm that genomic mutations present in fresh tumor were also detected in matched organoids (**Supplementary Fig. S2)**, consistent with prior studies (59).

**Figure 1.**
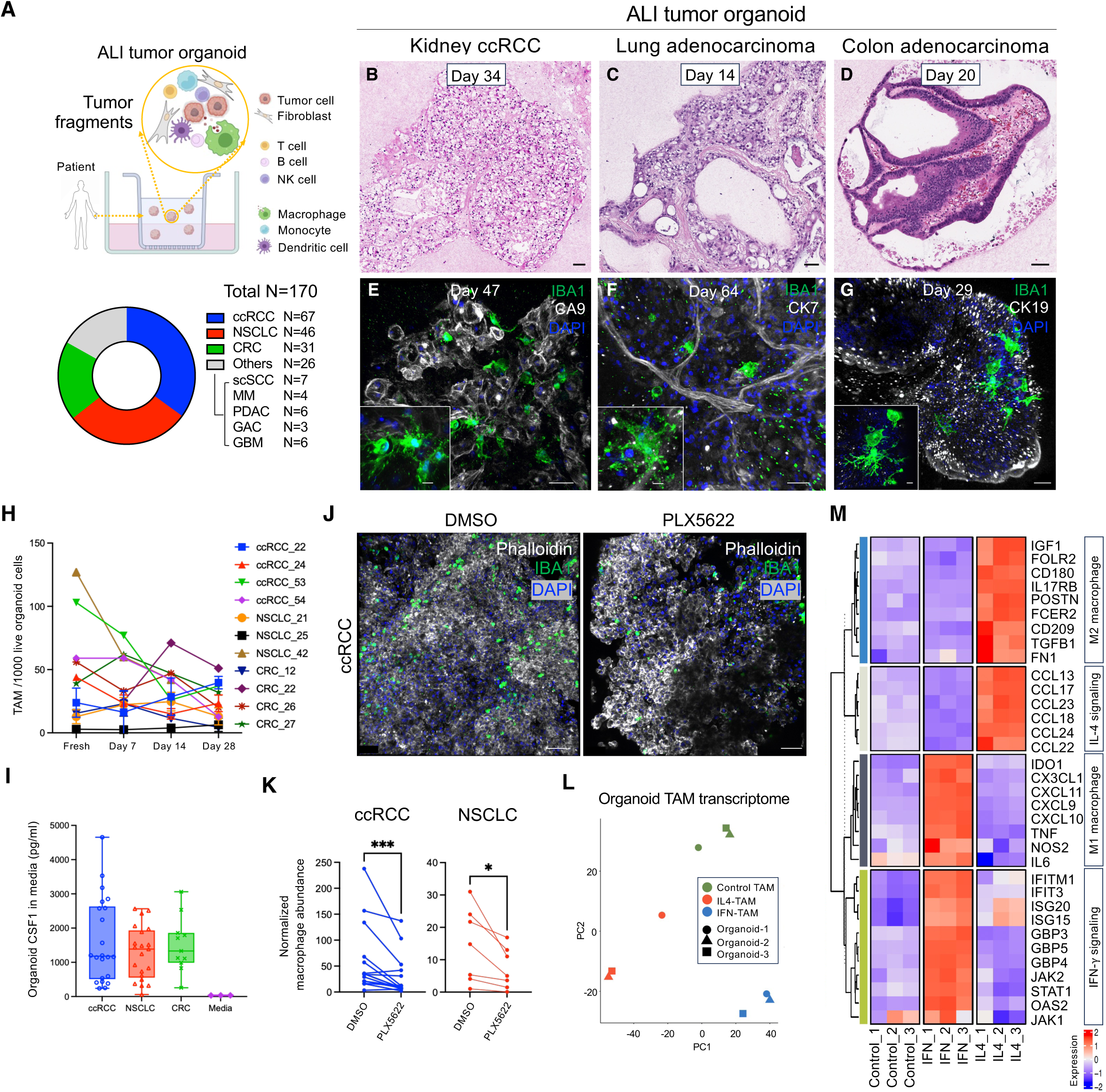
TAM within ALI patient-derived tumor organoids are maintained by endogenous CSF1 and exhibit functional responsiveness. **A**, Schematic depicting air-liquid interface (ALI) culture of patient-derived tumor organoids (PDO) from surgically resected tumor specimens and total biological replicates for all experiments aggregated by histologic subtype. **B-D,** Representative H&E staining of organoids from (**B**) clear cell renal cell carcinoma (ccRCC), culture day 34, (**C**) lung adenocarcinoma (LUAD), day 14 and (**D**) colorectal adenocarcinoma (CRC), day 20. Scale bars = 50 μm. **E-G,** Representative immunofluorescence staining of (**E**) ccRCC organoids, day 47, CA9 (white), IBA1 (green) and DAPI (blue), (**F**) LUAD organoids day 64, CK7 (white), IBA1 (green) and DAPI (blue), (**G**) CRC organoids day 29, CK19 (white), IBA1 (green), and DAPI (blue). Scale bars = 50 μm. **H,** Time course analysis of tumor-associated macrophage (TAM) abundance per 1000 live cells comparing fresh tumor versus matched day 7, 14, and 28 organoids (N=11 patients; 4 ccRCC, 3 NSCLC, 4 CRC). **I,** CSF-1 concentration in day 8 ALI organoid media measured by Luminex (N=54 patients; 22 ccRCC, 21 NSCLC, 11 CRC, 3 media only). Box plots represent mean and interquartile boundaries and whiskers extend to the minimum and maximum values. **J,** Representative immunofluorescence staining of day 8 ALI PDO (ccRCC) treated with DMSO or CSF-1R inhibitor PLX5622. IBA1 (green), phalloidin (white) and DAPI (blue), scale bar = 50 μm. **K,** Flow cytometry quantification of normalized macrophage abundance from (**J**) in day 8 ccRCC ALI organoids +/-CSF-1R-inihibitor PLX5622 for 8 days (N=23 patients, 15 ccRCC and 8 NSCLC), **=*p* <0.01, Wilcoxon test. **L,** Principal component analysis (PCA) plot of bulk RNA-seq of FACS-purified CD45+CD11b+HLA-DR+CD68+ TAM from day 8 ccRCC PDO after 24 h treatment with vehicle control, IFNγ- or IL-4, n=3 technical replicates. **M,** Heatmap of differential expressed genes (DEGs) from (**L**) depicting fold-induction of control-, IFNγ- and IL-4-TAM over averaged value of individual genes. Gene expression of IFN signaling, IL-4 signaling, M1-and M2-related gene signatures are depicted.

### Niche factor CSF-1 supports TAM survival in ALI organoids

The persistence of TAM without addition of exogenous macrophage/monocyte-related growth factors suggested that ALI organoid cultures intrinsically supported TAM survival. We performed Luminex cytokine analysis of organoid conditioned medium from different tumor histologies to evaluate secretion of colony-stimulating factor-1 (CSF-1), which is essential for macrophage/monocyte survival (11,64). The CSF-1 concentrations in ALI organoid conditioned medium at days 8-28 culture (longest time point attempted was day 70) were >1000 pg/ml across 3 tumor types (ccRCC, NSCLC, CRC), which are sufficient to maintain macrophage culture *in vitro* (65) (**Fig. 1I**) and vastly exceeded the CSF-1 present in media alone without cells (average 31.2 pg/ml) **(Supplementary Fig. S3A)**. We further evaluated if PDO

TAM survival required endogenously produced CSF-1 by treatment with the small molecule CSF-1R kinase inhibitor PLX5622, which is widely used for functional macrophage and microglia depletion (66). Accordingly, PLX5622 treatment decreased the prevalence of TAM in ccRCC and NSCLC organoids detected by flow cytometry (**Fig. 1J-K**). Further, PLX5622 treatment of GBM organoids converted microglia from an activated ameboid morphology to an inactive state with ramified processes (**Supplementary Fig. S3B-C**). These results suggest that the endogenous production of the essential macrophage growth factor CSF-1 maintains TAM abundance and activation state in ALI tumor organoids, and that TAM can be functionally manipulated in PDO by inhibiting the CSF-1 axis.

### Functional responsiveness of TAM within ALI tumor organoids

To further demonstrate the physiologic responsiveness of PDO TAM, we added known cytokines that polarize macrophages toward classical M1 and M2 phenotypes (67). Interferon-γ (IFNγ) was included in the culture media of day 8 tumor ALI organoids as a M1 phenotype inducer or IL-4 as a M2 phenotype inducer. After 24 hours treatment with IFNγ or IL-4, macrophages were purified from ALI organoid cultures by FACS. Bulk RNA-seq analysis revealed that the organoid TAM underwent distinct transcriptomic changes upon IFNγ versus IL-4 exposure (**Fig. 1L-M, Supplementary Fig. S4A-B, Supplementary Table S2**). Differentially expressed gene and pathway analysis comparing IFNγ and IL-4 exposures indicated that IFNγ-treated TAM (IFN-TAM) showed enrichment of IFN-inducible loci including *GBP5, JAK2, ISG15* and *IFITM1* (**Fig. 1M, Supplementary Fig. S4C**). IL-4 treated TAM (IL4-TAM) showed IL-4 related gene enrichment with higher expression of *CCL17, CCL18, CCL22* and *CCL24* (**Fig. 1M, Supplementary Fig. S4D**). IFN-TAM exhibited higher expression of the M1 macrophage-related genes *CXCL9/10/11, IDO1, TNF, IL6* and *NOS2 (***Fig. 1M, Supplementary Fig. S4E**) while IL4-TAM showed higher expression of the M2 macrophage-related genes *TGFB1, FN1, FCER2, IGF1, POSTN, CD209 and CD180* (**Fig. 1M and Supplementary Fig. S4F**). Immunofluorescence revealed TAM polarization toward M1 and M2 phenotypes in ALI organoids by CXCL9 and CCL17, respectively (**Supplementary Fig. S4G-I**). Upon flow cytometry, IFNγ consistently promoted expression of the M1 markers CD40 and CD80, while IL-4 induced the M2 marker CD206 in ALI organoid TAM cultures (**Supplementary Fig. S4J).**

### ALI tumor organoid TAM possess functional phagocytosis ability

We next assessed if organoid TAM exhibited phagocytic activity using imaging flow cytometry, in which high throughput cell analysis is combined with single cell microscopic imaging (68). TAM were isolated from ALI organoids (day 8-63) and then cultured with FITC-conjugated beads at 37°C or 4°C culture. Subsequent imaging flow cytometry allowed quantitation of phagocytic TAM that had functionally internalized beads by FITC positivity. The organoid TAM that had engulfed FITC bead conjugates were robustly captured at 37°C culture but were only infrequently observed at 4°C, indicative of specific phagocytic activity (**Fig. 2A-B**). Phagocytosis was further confirmed by measuring within intact ALI organoids the fluorescence intensity of the pH-sensitive fluorophore pHrodo, which is activated in low pH endosomes and lysosomes (69). Accordingly, organoid pHrodo fluorescence was prominently detected in PDO and was partially reversed by cytochalasin D which inhibits actin polymerization necessary for efficient phagocytosis, indicative of a specific signal (**Fig. 2C**). Further, we confirmed phagocytosis within TAM by the overlay of pHrodo and CD11b+ cells within intact organoids (**Fig. 2D-E**). Overall, TAM phagocytic capacity was well preserved in ALI tumor organoid culture (**Fig. 2A-E**).

**Figure 2.**
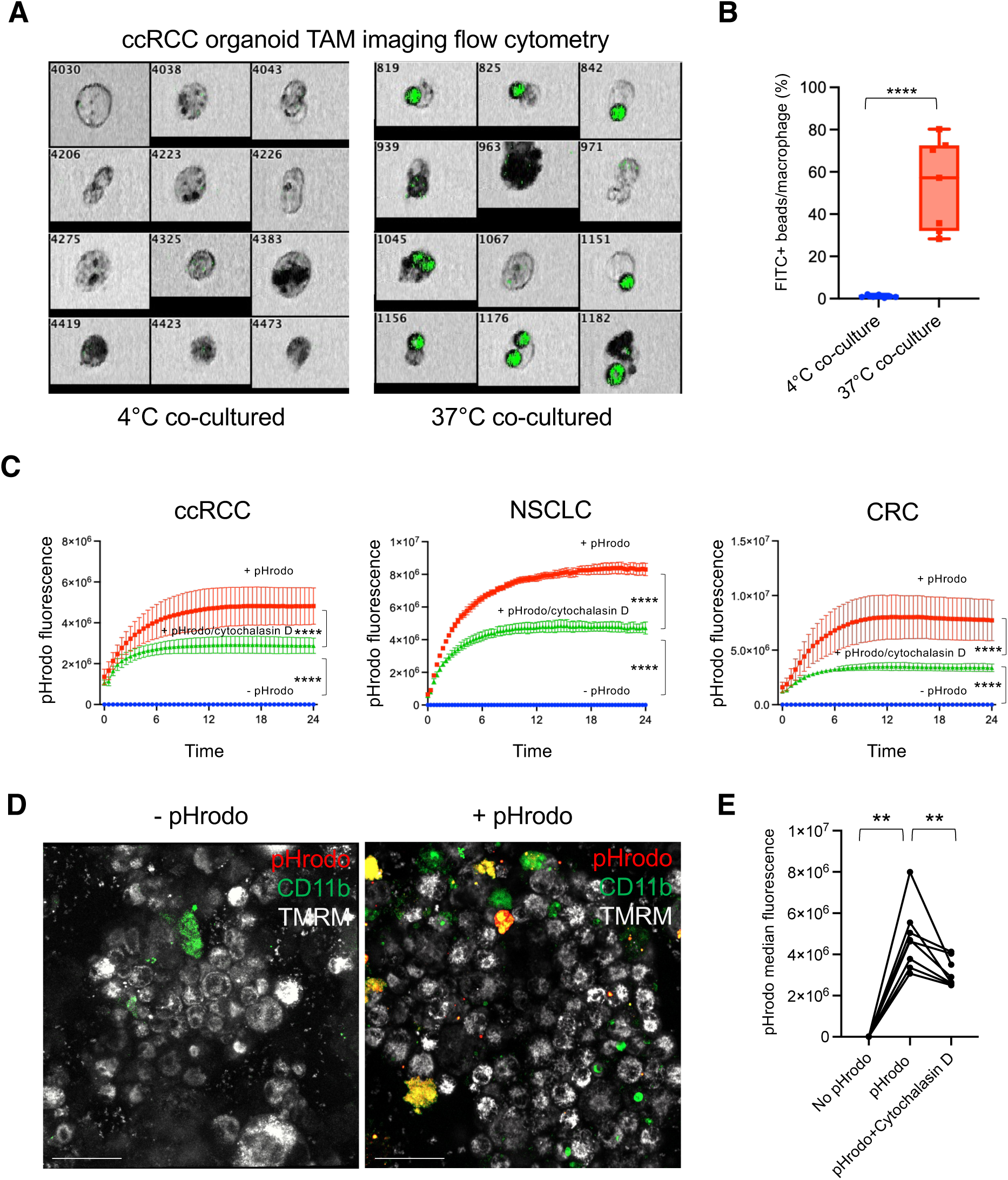
Organoid TAM conduct functional phagocytosis. **A**, Representative image of fluorescent bead phagocytosis assay using imaging flow cytometry (IFC), gated on macrophages (organoid TAM). Single-cell suspensions of NSCLC day 10 ALI organoids were co-cultured with FITC-labeled beads at 4 ℃ or 37℃ and single CD45+CD11b+HLA-DR+CD68+ macrophages were imaged by IFC for the presence of phagocytosed beads (green). **B,** Quantification of (**A**) showing frequencies of FITC+ bead positivity in organoid TAM (day 8-45) (N=7 patients), ****=*p* <0.0001, Wilcoxon test. Box plots represent mean and interquartile boundaries and whiskers extend to the minimum and maximum values. **C,** Representative real time kinetics of phagocytosis assay showing pHrodo bioparticle uptake in ccRCC, NSCLC and CRC organoids (day 10-63) with or without cytochalasin D treatment. **D,** Representative live cell organoid imaging from (**C**) showing pHrodo-positive cells co-localized with CD11b+ cells. Data are represented as mean ± SD. (**E**) Quantification of (**C**) showing organoid median pHrodo fluorescence intensity at 12 hours (N=8 patients), **=*p* <0.01, Mann-Whitney test.

### Organoid TAM tumor cell phagocytosis enhancement by anti-CD47

We examined the modulation of tumor phagocytosis in ALI organoids by employing anti-CD47 as proof-of-principle for targeting negative inhibitory checkpoints mediated by anti-phagocytic “don’t eat me” signals. We confirmed that the anti-CD47 antibody B6H12, which functionally inhibits the CD47-SIRPα interaction (35,41,42), achieved quantitative CD47 occupancy in organoid tumor cells by pronounced inhibition of an anti-CD47 flow cytometry detection antibody (**Supplementary Fig. S5A**). The B6H12 clone was thus used for all anti-CD47 experiments in subsequent studies. TAM were isolated by anti-CD11b bead enrichment from ALI organoids (ccRCC, NSCLC, CRC) and labeled with CellTrace^TM^ Far Red. In parallel, ALI organoid tumor epithelium from identical patients was isolated by anti-EpCAM beads and labeled with Calcein AM. The isolated CellTracker^TM^ Deep Red-labeled organoid TAM were then combined with autologous Calcein AM-labeled tumor epithelium with and without the B6H12 anti-CD47 antibody. In this phagocytosis assay, anti-CD47 increased the uptake of Calcein AM-labeled fluorescent tumor cells within CellTrace^TM^ Far Red dye-positive ALI tumor organoid-derived TAMs as determined by CD11b+ Far Red+ Calcein AM+ events (**Fig. 3A-B**). This indicated that tumor phagocytosis by ALI PDO TAM can be promoted by inhibition of the CD47 anti-phagocytic signal.

**Figure 3.**
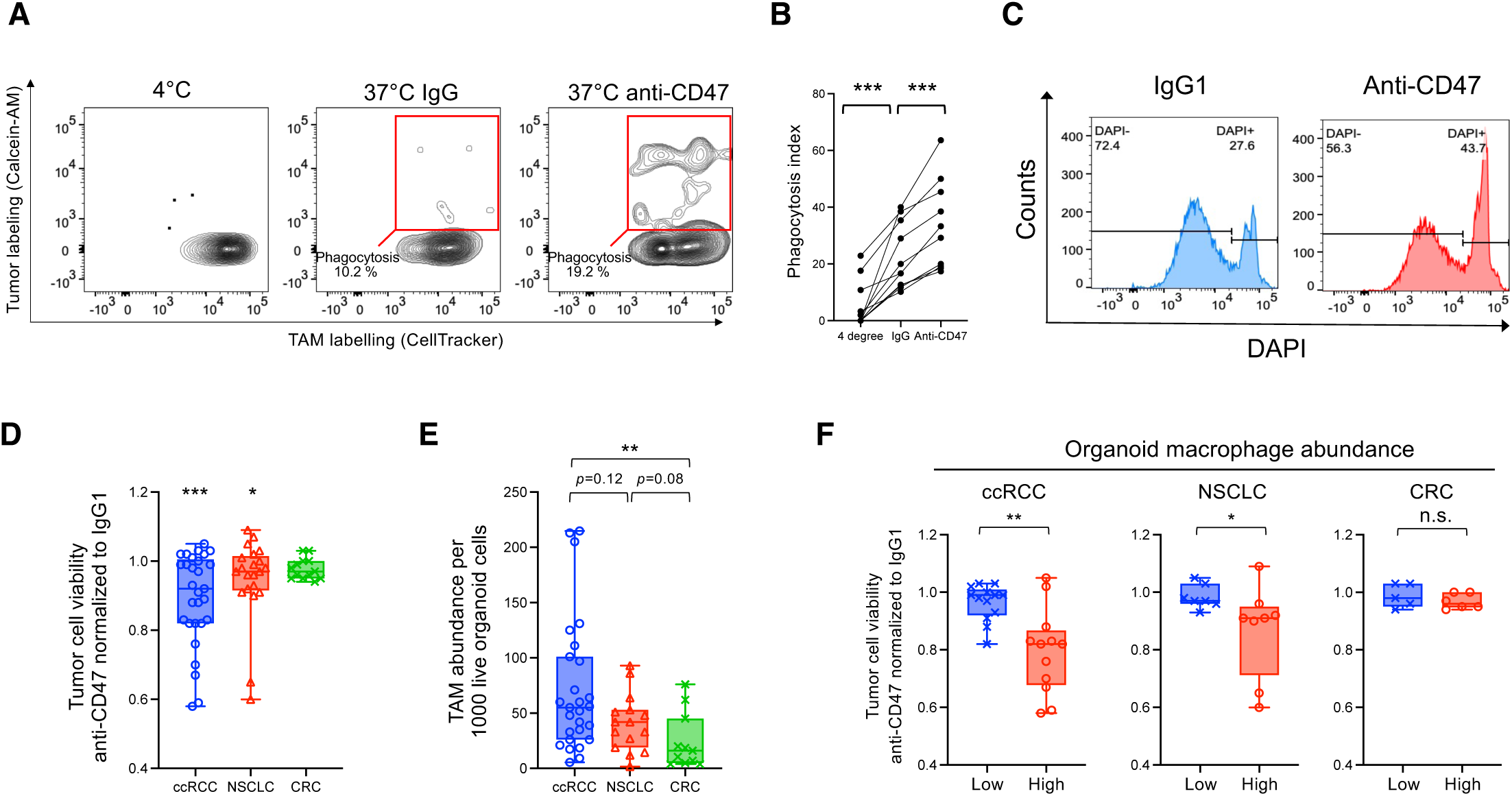
Organoid screening of anti-CD47-responsive tumor histologies. **A**, Representative flow cytometry plots of organoid TAM tumor phagocytosis assays. Matched organoid TAM and tumor epithelium from ALI PDO were isolated by magnetic beads, labeled with Calcein-AM and CellTracker respectively, and then co-cultured. Tumor phagocytosis of Calcein-AM+ tumor cells by Celltracker+ organoid TAM was indicated by double-positive cells within the red box. **B,** Quantification of (**A**) (N=11, 5 ccRCC, 2 NSCLC and 4 CRC). **C,** Representative flow cytometry histogram of CD45-EpCAM+ pre-gated live (DAPI-) versus dead (DAPI+) organoid tumor cells in CRC organoids after anti-CD47 or IgG1 treatment, day 8. **D,** Box plots of fractional tumor cell viability in anti-CD47-treated day 8 ALI organoids determined by flow cytometry and normalized to IgG1 control as in (**C**). N=62; 27 ccRCC, 22 NSCLC, 13 CRC. *=*p*<0.05, ***=*p*<0.001 versus IgG1, Wilcoxon test. **E,** CD45+CD11b+HLA-DR+CD163+ macrophage abundance per 1000 live organoid cells from (**D**) quantified by flow cytometry (N=51 patients; 25 ccRCC, 15 NSCLC, 11 CRC). **F,** Fractional tumor cell viability of TAM-low versus TAM-high ALI PDO from (**A**), ccRCC: low N=13, high N=12, NSCLC: low N=7, high N=8, CRC: low N=5, high N=6. A median value for macrophage abundance per 1000 live organoid cells (ccRCC: 55.5, NSCLC: 42, CRC: 16) was used as a threshold for TAM-low versus TAM-high status. **=*p*<0.01, *=*p*<0.05 Mann-Whitney test.

### Organoid screening of anti-CD47 responsive tumor histologies

To explore if PDO could screen tumors to identify histologic subsets that respond to a given therapy, thus informing clinical trial design and patient selection, we therefore used ALI organoids to identify potential anti-CD47-responsive tumor types. PDO were screened for anti-CD47-induced tumor killing across 8 tumor histologies from a total of 82 patients, including ccRCC (N=27), NSCLC (N=22), CRC (N=13) and other tumors (N=20). The cytotoxic effects of anti-CD47 organoid treatment were measured after 8 days by the decreased fractional viability of CD45-EpCAM+ organoid tumor epithelium using flow cytometry with fixable cell viability dyes (**Fig. 3C**). Anti-CD47 promoted organoid tumor epithelial cell death most prominently in ccRCC as opposed to NSCLC and CRC **(Fig. 3D)**; effects in other histologies were not observed but cannot be excluded as they were underrepresented in our screening panel (**Supplementary Fig. S5C**). Tumor fragments could be detected within TAM from intact organoids upon anti-CD47 treatment (**Supplementary Fig. S5B**). Notably, TAM tended to be more abundant in organoids from ccRCC, which was significant versus CRC but not NSCLC or other tumor types (**Fig. 3E and Supplementary Fig. S5D**). The high TAM content of ccRCC organoids is consistent with CD68 enrichment in ccRCC versus diverse tumor histologies in pan-cancer TCGA data (**Supplementary Fig. S 5E**) and in several published studies (12,70–72). The degree of TAM infiltration stratified ccRCC organoid responders versus non-responders, where higher TAM organoid content was associated with increased anti-CD47-induced organoid tumor epithelial killing **(Fig. 3F)**. NSCLC organoids, which displayed a generally weaker anti-CD47 cytotoxic response **(Fig. 3D)**, also exhibited correlation of tumor killing with TAM content, while CRC organoids could not be stratified in this manner (**Fig. 3F**).

### Concordance of secreted cytokines between anti-CD47 treated organoids and patients

To more broadly evaluate TAM cytokine production in PDO, Luminex cytokine array analysis was performed on culture supernatants from anti-CD47-treated day 8 tumor organoids from 49 patients (19 ccRCC, 20 NSCLC, 10 CRC). Higher secretion of numerous cytokines related to pro-inflammatory macrophages (CCL3, CCL4 and TNFA) were detected in the conditioned medium of organoids treated with anti-CD47-treated compared with IgG control, while IFNγ, CXCL9 and CXCL10 were not elevated (**Fig. 4A and Supplementary Table S3**).

**Figure 4.**
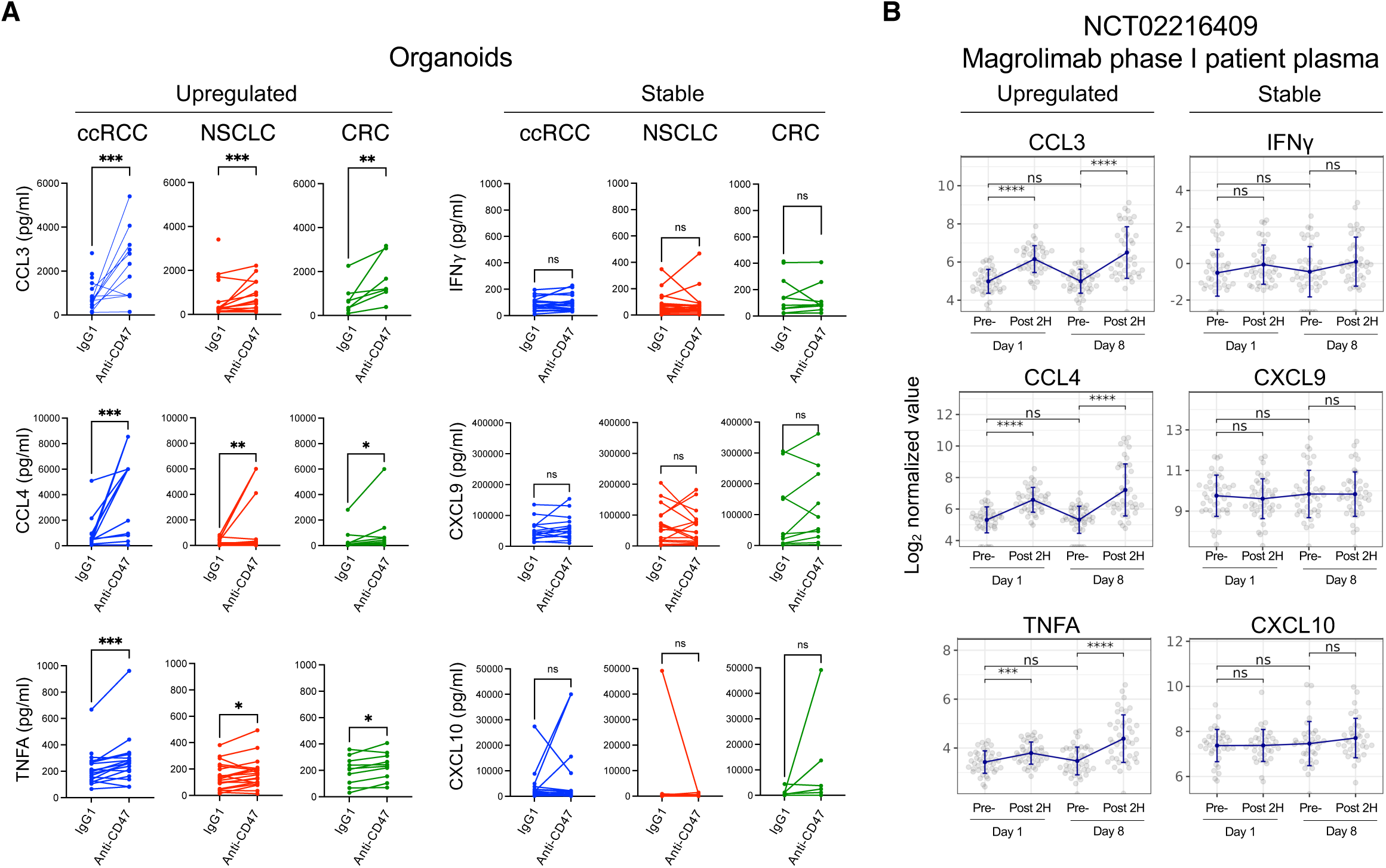
Concordance of secreted cytokine biomarkers between anti-CD47-treated organoids and patients. **A**, Luminex quantification of cytokine release into the conditioned media in IgG1- or anti-CD47-treated day 8 ALI organoids. The supernatant was collected on day 8 culture, with the last media change 3 days prior. The IgG1 and anti-CD47 conditions were harvested in parallel for each biological replicate pair. Samples exceeding the linear range of detection were excluded from the analysis (**Supplementary Table S3**). CCL3 (15 ccRCC, 18 NSCLC, 8 CRC), CCL4 (14 ccRCC, 17 NSCLC, 9 CRC), TNFA (19 ccRCC, 20 NSCLC, 10 CRC), IFN**γ** (19 ccRCC, 20 NSCLC, 10 CRC), CXCL9 (15 ccRCC, 20 NSCLC, 10 CRC) and CXCL10 (16 ccRCC, 17 NSCLC, 6 CRC). *=*p*<0.05, **=*p*<0.01, ***=*p*<0.001, Wilcoxon test. **B,** Quantification of cytokines from patient plasma in an anti-CD47 magrolimab monotherapy phase I clinical trial (NCT02216409)(40). Cytokines were measured using fit-for-purpose validated Simple Plex^TM^ assays (Bio-Techne, Minneapolis, MN). Plasma was collected before magrolimab infusion (“pre”) and 2 hours after infusion (“post”) at day 1 (priming dose 1 mg/kg) and day 8 (loading dose 20-45 mg/kg) from 41 advanced solid tumor patients. No cytokine levels were above ULoQ. For measurements below LLoQ, we imputed values with an offset of 0.1. *=*p* <0.05, \*\**=p* <0.01, ***=*p* <0.001, ****=*p* <0.0001; Mann-Whitney test. ULoQ and LLoQ refer to upper and lower limits of quantitation, respectively.

We confirmed these organoid observations in patient populations by analyzing plasma from a phase I clinical trial of patients with advanced solid tumors treated with the anti-CD47 antibody magrolimab (NCT02216409) (40). In this trial, magrolimab was administered at 1 mg/kg (priming dose) on study day 1 and at 20-45 mg/kg (loading dose) on day 8, followed by multiplexed cytokine plasma immunoassay. Plasma CCL3, CCL4 and TNFA were elevated after magrolimab priming (day 1) and loading (day 8) doses versus pre-administration samples from the same patients **(Fig. 4B)**. Notably, this paralleled elevations of CCL3, CCL4 and TNFA in anti-CD47-treated organoids **(Fig. 4A)**. Importantly, cytokines related to macrophage functions such as IFNγ, CXCL9 and CXCL10 that were not elevated by anti-CD47 in organoids, similarly were not upregulated in the magrolimab-treated clinical cohort (**Fig. 4A-B**). Taken together, anti-CD47 cytokine endpoints from ALI tumor organoids were concordant with pharmacodynamic measurements in patients.

### Anti-CD47 induces dynamic changes in TAM and promotes the SPP1+ phenotype

To investigate anti-CD47-regulated TAM phenotypic changes in single cell resolution, we next performed scRNA-seq of the CD45+ hematopoietic fraction organoid cultures from tumors of 10 distinct patients (6 ccRCC and 4 NSCLC) after 8 days of treatment with anti-CD47- or IgG1. In parallel, scRNA-seq was performed on matched fresh tumor for 5 of the ccRCC and 2 of the NSCLC organoid cultures. ALI PDO maintained essential immune components including myeloids, mast cells, CD4 and CD8 T cells, B cells and NK cells (**Fig. 5A and Supplementary Fig. S6A-D**).

**Figure 5.**
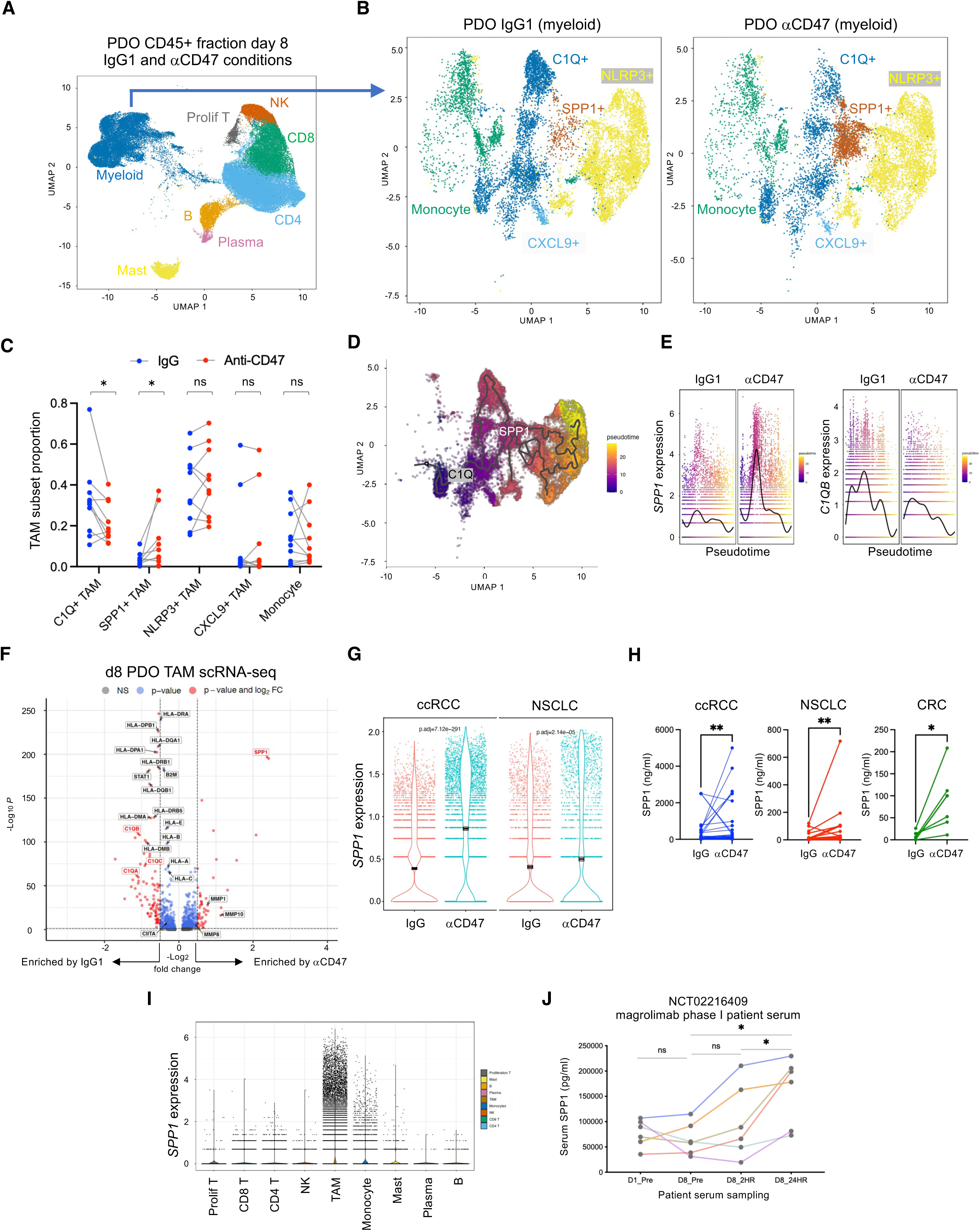
Anti-CD47 induces dynamic changes in organoid TAM and promotes the SPP1+ phenotype. **A**, UMAP plot of single cell RNA-seq from the FACS-sorted CD45+ fraction of 10 ALI PDO (6 ccRCC and 4 NSCLC) cultured at day 8 showing major immune subsets. **B,** High-resolution UMAP subclustering of the scRNA-seq TAM and monocyte clusters from (**A**) depicting PDO TAM subset differences in IgG1-versus anti-CD47-treated day 8 ALI organoids, N=10 patients (6 ccRCC, 4 NSCLC). **C,** Quantification of (**B**) depicting proportion of TAM subsets per total TAM,^*^=*P*<0.01, Mann-Whitney test. **D,** Monocle3 trajectory analysis of TAMs in PDO from (**B**) excluding monocytes. Approximate locations of cell types are labeled. **e,** *SPP1* and *C1QA* expression changes along the pseudotime axis from (**D**). **F,** Volcano plot of differentially expressed genes between IgG1- or anti-CD47-treated PDO TAM, culture day 8, N=10 patients (6 ccRCC, 4 NSCLC). Log_2_ fold change is shown on the x axis and -log_10_ adjusted *p* value on the y axis. *p* value of 0.05 and fold change of 1 are indicated. **G,** *SPP1* gene expression induction by anti-CD47 in organoid TAM from ccRCC (N=6) and in NSCLC (N=4) as in (**F**). **H,** SPP1 ELISA of conditioned media from IgG- or anti-CD47-treated day 8 ALI organoids (N=59 patients; 33 ccRCC, 19 NSCLC and 7 CRC). *=*p* <0.05, ** = *p*<0.01, Wilcoxon test. **I,** *SPP1* gene expression in UMAP plots of PDO CD45+ cells from (**A**). **J,** Quantification of secreted SPP1 from patient serum in an anti-CD47 magrolimab monotherapy phase I clinical trial (NCT02216409)(40) measured by Luminex (N=6 patients). Serum was collected before infusion and 2 hours or 24 hours after magrolimab infusion at day 1 and day 8. *=*p* <0.05, Wilcoxon test.

Recent single cell analyses have revealed substantial phenotypic diversity within TAM, including *SPP1+*, *NLRP3+*, *C1Q+* and *CXCL9+* subsets (12–16). Accordingly, unsupervised subclustering of the myeloid scRNA-seq populations from **Fig. 5A** revealed prominent *SPP1+*, *NLRP3+*, *C1Q+* and *CXCL9+* TAM, and monocyte clusters **(Fig. 5B and Supplementary Fig. S6E-H**). TAM subsets were well conserved in day 8 ALI PDO compared with fresh tumor, but revealed an expected decrease in monocytes, which exhibit a lifespan of 1-7 days (63) (**Supplementary Fig. S6E-H)**. The main TAM subsets (*C1Q*+, *SPP1*+, *NLRP3*+, *CXCL9*+) were conserved in both organoids and cognate fresh tumors but organoids exhibited a disproportionate increase in *NLRP3+* TAM (**Supplementary Fig. S6E**).

Anti-CD47 notably stimulated the abundance of organoid SPP1+ TAM, which have been linked to immunosuppression (21–26). Simultaneously, anti-CD47 repressed organoid C1Q+ TAM (**Fig. 5B-C)**. Notably, *SPP1+* TAM stimulation by anti-CD47 was greater in ccRCC than in NSCLC (**Fig. 5E and Supplementary Fig. S7A-B**). Pseudotime trajectory analysis suggested that CD47 inhibition promoted a cell state progression where C1Q+ TAM phenotypically converted into SPP1+ TAM (**Fig. 5D-E**), inferring that the CD47 pathway physiologically regulates interconversion between these two TAM subsets. Consistent with a cell state transition, anti-CD47 did not alter proliferative transcripts in organoid TAM **(Supplementary Fig. S7C)**.

Anti-CD47 induced transcripts for numerous cytokines, chemokines and secreted factors within organoid TAM, including *SPP1*, *MMP1, MMP9* and *MMP10* related to fibrosis and tissue remodeling(13,14) (**Fig. 5F, Supplementary Fig. S7D and Supplementary Table S4**). Interestingly, anti-CD47 downregulated TAM gene expression profiles including transcripts for C1Q+ TAM (*C1QA, C1QB, C1QC),* regulators of MHC class II expression (*CIITA)*, MHC class-I antigen presentation pathway (*B2M, TAP1, TAP2),* MHC class -I and -II (*HLA-DRA, HLA-DPB1*, *HLA-DQA1, HLA-A, HLA-B, HLA-C)*, and interferon response (*ISG15, GBP1, IRF1, STAT1, IFITM3, CXCL10)* (**Fig. 5C and Supplementary Fig. S7E**). These transcriptomic changes were observed both in ccRCC and NSCLC organoid TAM (**Supplementary Fig. S7D-E**). Concordantly, pathway analysis revealed that the strongest processes downregulated in anti-CD47-treated organoid TAM included MHC-II antigen presentation, MHC-I cross-presentation and IFN signaling (**Supplementary Fig. S7F**).

Notably, *SPP1* was the most strongly anti-CD47-induced mRNA in organoid TAM and was paralleled by robust downregulation of *C1QA, C1QB* and *C1Q* mRNA (**Fig. 5F**). Further, *SPP1* stimulation by anti-CD47 was greater in ccRCC than in NSCLC (**Fig. 5G and Supplementary Fig. 7A-B**). Crucially, the elevation of *SPP1* mRNA was concordant with the anti-CD47 promotion of the *SPP1+* TAM subset (**Fig. 5B-C**). In parallel with increased *SPP1+* TAM, CD47 inhibition promoted SPP1 secretion in ELISA assays of organoid culture supernatants from multiple cancer types (**Fig. 5H**). Further, *SPP1* expression was specific for TAM but not other immune cell populations upon scRNA-seq **(Fig. 5I)**.

The possibility that anti-CD47-stimulated SPP1 production also occurred in human populations was explored in serum from the NCT02216409 phase I magrolimab clinical trial (40). Strikingly, magrolimab administration to patients at day 8 elevated SPP1 levels in patient serum after 2 hours with further increases after 24 hours (**Fig. 5J**). In summary, the concordant SPP1 elevations induced by anti-CD47 in both organoids and patients reiterate that organoids faithfully recapitulate in vivo human TAM biology across tumor histologies, and reveal induction of the immunosuppressive SPP1+ TAM subset as an unforeseen complication of CD47 blockade.

## Discussion

TAM exert manifold phagocytic, paracrine and antigen-presenting functions within the TME(7–9), which have raised significant interest in their therapeutic targeting (4–6). However, experimental systems to study endogenous TAM within their native human TME have been notably lacking, despite extensive reconstitutive studies in which tumor cells are co-cultured with exogenous myeloid elements (73,74). We previously described PDO cultures in which primary human tumor fragments cultured in an ALI successfully retained a diverse immune TME spanning T, B, NK and myeloid subsets without requiring artificial reconstitution (59). Here, we extended these studies to deeply characterize the TAM content of ALI PDO. The organoid TAM were maintained for >70 days as supported by endogenous CSF-1 production and were functionally competent in phagocytosis, polarization induction and cytokine secretion. Major TAM subsets that have been as extensively described by single cell RNA-sequencing(12–16) (*C1Q+, SPP1+, NLRP3+, CXCL9+*) were conserved in both organoids and cognate fresh tumors. Interestingly, organoids exhibited substantially increased *NLRP3+* TAM versus fresh tumors, which could reflect *in vitro* culture conditions.

As proof-of-principle for functional TAM exploration in the ALI organoid system, anti-CD47 blocking antibodies stimulated tumor phagocytosis but also enabled mechanistic identification of events regulated by the CD47-SIRPα pathway. Unexpectedly, anti-CD47 stimulated an increase in *SPP1+* TAM, accompanied by decreased *C1Q+* TAM. Our results support a model where CD47 inhibition directly induces a cell fate transition of *C1Q+* TAM into *SPP1+* TAM, based upon scRNA-seq pseudotime analysis and lack of anti-CD47 proliferative effects on *SPP1+* or *C1Q+* TAM. Further, these findings are aligned with the known mutual exclusivity of *SPP1+* and *C1Q+* TAM (30,31). Conversely, this infers that the CD47 pathway physiologically regulates TAM subset identity, potentially by promoting the *C1Q+* TAM cell fate at the expense of *SPP1+* TAM.

Notably, CD47 inhibition stimulated two distinct negative feedback mechanisms that could oppose tumor immunity. Firstly, the induction of *SPP1+* TAM could obligately dampen immune responses after phagocytosis, and actively oppose beneficial anti-tumor activities of CD47 inhibition. Indeed, in addition to facilitating matrix remodeling and fibrosis (13,17,18), *SPP1+* TAM exhibit immunosuppressive functions (21–24) and *SPP1+* TAM tumor infiltration is related to poor prognosis (19) and dysfunctional and regulatory T cells (75). Further, *SPP1+* TAM mediate immune checkpoint therapy resistance (25,26) through adenosine signaling, emphasizing their importance as a therapeutic target and biomarker for treatment response (25). SPP1 interaction with its receptor CD44, and spatial association of *SPP1+* TAM and *CD44+* cancer cells are associated with treatment resistance and cancer stemness (76,77). The anti-CD47 induction of *SPP1+* TAM could be either dependent or independent of phagocytosis, given the numerous downstream effector functions of the CD47 pathway (37–39). Secondly, anti-CD47 also downregulated MHC in organoid TAM, potentially reflecting in part the decreased abundance of MHC class II-expressing *C1Q+* TAM (27,28). The resultant decrease in antigen presentation capacity could further exacerbate the anti-CD47-induced immunosuppressive and TME remodeling effects of *SPP1+* TAM. Thus, CD47-SIRPα pathway blockade promotes phagocytosis by TAM but obligately induces two distinct negative feedback mechanisms that could underlie the lack of clinical efficacy of current CD47 inhibitors (45,46). These results further suggest rational strategies for improving the antitumor efficacy of blocking CD47 or of other TAM regulatory checkpoints.

Our data also demonstrate the potential of organoids to identify responsive tumor histologies for macrophage-targeted therapeutics. Amongst the ccRCC, NSCLC and CRC tumor ALI cultures screened, ccRCC organoids manifested the highest degree of anti-CD47-induced tumor killing. This ccRCC organoid susceptibility was linked to a higher degree of TAM infiltration, consistent with reports of TAM enrichment in ccRCC tumors amongst diverse cancer histologies (12,70–72). In contrast, anti-CD47 promoted SPP1 secretion and CCL3/CCL4/TNFA cytokine release across all tumor histologies examined, which may represent more promiscuous responses to CD47 inhibition that are not sufficient to confer anti-CD47 tumor cytotoxicity, which was strongest in ccRCC organoids. In parallel with scRNA-seq, anti-CD47 increased SPP1 secretion in both organoid culture supernatant and in patient serum, although this could reflect either *SPP1+* TAM expansion and/or increased *SPP1* transcription. Of note, increased circulating SPP1 levels are associated with more advanced disease and tumor progression in cancer (78–81). Our findings also suggest that elevated SPP1 secretion comprises a robust biomarker for anti-CD47 treatment that could further mediate tumor therapeutic resistance.

The ccRCC organoid response to proof-of-principle CD47 blockade also suggests the potential benefit of targeting this tumor type with next-generation macrophage checkpoint inhibitors, such as anti-CD47 antibodies with disabled Fc regions (47–49) or targeting SIGLEC-10 and LILRB1 (36,82). This will require clinical validation, as macrophage checkpoint inhibition has not yet been evaluated in ccRCC trials. Notably, any potential anti-tumor activity of inhibition of SIGLEC-10 or LILRB1 could potentially be obligately blunted by identical or similar negative feedback mechanisms as for CD47/SIRPα in the present studies. Here, the anti-CD47 tool antibody blocked the SIRPα inhibitory signal and provided a pro-phagocytic signal through Fc. The addition of tumor antigen-targeting antibodies and Fc domain engineering (83) could further enhance this activity and specificity against cancer histologies beyond ccRCC but could elicit different downstream responses from the present study. PDO may also nominate clinical biomarkers, since the observed anti-CD47 induction of cytokine secretion (CCL3, CCL4, TNFA) and SPP1 in organoids were all confirmed in patient plasma and serum from a magrolimab monotherapy phase I trial (40).

Overall, the present TME organoid model should enable mechanistic investigations of tumor macrophage biology, immune network interactions and treatment resistance that have been previously inaccessible to conventional *in vitro* methods. The intact human immune TME of ALI organoids may also allow biomarker discovery and precision medicine identification of tumor types and patient subsets that might optimally respond to functional macrophage modulation, such as ccRCC. Such studies could also facilitate pre-clinical development of next-generation TAM therapeutics for targets beyond our proof-of-principle explorations with anti-CD47. Finally, through recapitulation of the *in vivo* biology of macrophages and their interactions with other immune subsets and epithelium, the current organoid method may find application to additional categories of human disease, such as infectious disease and autoimmunity.

## Methods

### Human specimens

Tumor specimens from surgical resections used for ALI patient-derived tumor organoid (PDO) generation were obtained through the Stanford Hospital Tissue Bank. All experiments using human tissue were approved by the Stanford University Institutional Review Board. Tumors from 170 patients included clear cell renal cell carcinoma (ccRCC; N=67), non-small cell lung cancer (NSCLC; N=46), colorectal adenocarcinoma (CRC, N=31), cutaneous squamous carcinoma (cSCC; N=7), malignant melanoma (MM: N=4), pancreatic adenocarcinoma (PDAC; N=6), gastric adenocarcinoma (GAC: N=3), and glioblastoma GBM; (N=6) under IRB-28908 approved by the Research Compliance Office of Stanford University. Patient characteristics are presented in **Supplementary Table S1**.

### Establishment of ALI tumor organoid cultures

Transwell inserts (60 mm^2^) with a membranous bottom (Millipore, PICM03050) were placed into wells of a 6-well plate. Collagen mixtures were prepared by mixing Cellmatrix Type I-A (FUJIFILM Wako,637-00653), 10x concentrated Ham’s F-12 (Gibco, 21700075), and reconstitution buffer (260 mM NaHCO_3_, 0.05 M NaOH and 200 mM HEPES) on ice at a ratio of 8:1:1. 1 ml of reconstituted collagen mixture was added to each insert, which served as a bottom layer without tissue. This bottom layer was left to solidify for 10 min in a 37°C incubator. Surgically resected tumor tissues were minced finely with iris scissors on a petri dish and added to the remaining collagen mixture after wash with PBS. 1 ml of WENR media (50% ADMEM/F12 (Gibco, 12634028) with 50% WNT3A-, RSPO1-, Noggin-containing media (L-WRN, CRL-3276^TM^, ATCC) and HEPES (1 mM, Gibco, 15630080), Glutamax (1X, Gibco, 35050061), nicotinamide (10 nM, Sigma, N0636-500G), N-acetylcysteine (1 mM, Sigma, A9165-100G), B-27 without vitamin A (1X, Gibco, 125870-01), A83-01 (0.5 μM, Tocris, 2939), Pen-Strep glutamine (1X, Gibco, 10378016), gastrin (10 μM, Sigma, G9145), EGF (50 ng/ml, PeproTech, AF-100-15)), and supplemented with Normocin (Invitrogen, ant-nr-2), 5% FBS (Biotechne, S11550), 10 μM Y-27632 dihydrochloride (Biogems, 1293823), 10 μM CHIR 99021 (Biogems, 2520691), IL-2 (100 IU/ml, Peprotech, 200-02) was added into each well of the 6-well plate outside of the inserts, generating an air-liquid interface (58,59).

### ALI tumor organoid culture and immune modulation treatments

All experiments modulating the ALI organoid immune component were performed in WENR media described above. To block the CD47-SIRPα axis, an anti-human CD47 antibody (BioXcell, clone: B6.H12, BE0019-1) or human IgG1 isotype control (BioXcell, BE0297) were added to the culture medium, each at 20 μg/ml, for 7 days. For macrophage polarization experiments, recombinant human IFN-γ (PeproTech, 300-02) was added at 100 ng/ml for M1 polarization and recombinant human IL-4 (PeproTech, 200-04) at 50 ng/ml for M2 polarization. After 24 hours incubation, ALI tumor organoids were analyzed and macrophages were isolated by FACS. For macrophage loss-of-function experiments, PLX5622 (Selleckchem, S8874) was used at 10 nM and analyzed at day 8 by flow cytometry and IF staining.

### Organoid cryopreservation

ALI PDO were generated as above and grown for at least 5 days. Each organoid containing collagen gel was carefully removed from the insert and replaced into 1 ml of cryopreservation media (Bambanker) and stored at -80℃ for at least 24 h. For cryorecovery, gels were quickly thawed in 37℃ water bath and each organoid-containing collagen gels were washed with WENR media for 2 times in tissue culture dishes. Gels were then mounted on the top of bottom layer of collagen gel as described, and a further 500 µl of collagen-mixture was layered on top. After solidifying for 10 min in a 37℃ incubator, 1 ml of media was added to each well.

### Immunofluorescence

Unstained sections from FFPE were deparaffinized and rehydrated, followed by antigen retrieval using citrate-based buffer (MilliporeSigma, C9999) at 95℃ for 20 minutes in a steamer. Sections were blocked with 10% donkey serum (Jackson ImmunoResearch, 017-000-121) for 1 hour at room temperature (RT). Primary antibodies were incubated overnight at 4℃, followed by secondary antibodies incubation with DAPI for 1 hour at RT. Sections were mounted in mounting medium (Vector Laboratories, H-5501) and cover-slipped. As for whole mount immunofluorescence, the organoid containing collagen matrix was cut into smaller pieces, then fixed in 4% paraformaldehyde (PFA) for 1 h at RT and washed with PBS. PFA was quenched with PBS-glycine (130 mM NaCl, 13.2 mM Na_2_HPO_4_, 3.5 mM NaH_2_PO_4_, 100 mM glycine in PBS at pH 7.4) for 30 minutes at RT with gentle rocking. After washing, the collagen pieces were incubated in blocking solution (10% donkey serum diluted in permeabilizing solution (130 mM NaCl, 13.2 mM Na_2_HPO_4_, 3.5 mM NaH_2_PO_4_, 7.7 mM NaN_3_, 15 μM BSA, 2% Triton X-100, 0.5% TWEEN-20 in PBS at pH 7.4) for 2 hours at RT with gentle rocking. The collagen pieces were then stained with primary antibodies diluted in blocking solution for 1-3 days at RT with gentle rocking. After washes with permeabilizing solution, the collagen pieces were incubated with secondary antibodies diluted in blocking solution for 4 hours at RT with gentle rocking. The collagen pieces were transferred and mounted on slide. Vacuum grease was used around the collagen pieces to avoid flattening the organoids inside. Alternatively, whole mount staining was performed by dissolving collagen gels with collagenase IV (Worthington, LS004212) at 37°C for 40 min to release intact organoids followed by the same procedure as above. All images were taken using Keyence BZ-X700 microscope or Zeiss LSM900 or LSM980 confocal microscopes. Immunofluorescence staining of sections and whole-mount organoids used the following anti-human antibodies: anti-CD68 (Abcam, ab213363), anti-CD68 (Agilent Dako, M0876), Alexa Fluor 555-anti-CD68 (Abcam, ab279323), anti-IBA1 (FUJIFILM Wako, 019-19741), anti-PAX8 (Proteintech, 10336-1-AP), anti-TTF1 (Abcam, ab76013), anti-pan-cytokeratin (Abcam, ab7753), Alexa Fluor 488-anti-pan-cytokeratin (Abcam, ab277270), anti-cytokeratin 19 (Biotechne, AF3506), anti-EpCAM (Abcam, ab79079), anti-EpCAM (Abcam, ab223582), anti-GFAP (Thermo Fisher Scientific, 13-0300), anti-melanoma (Abcam, ab732), anti-CXCL9 (Biotechne, AF392), anti-CCL17 (Abcam, ab195044), anti-CD3 (Abcam, ab11089) and anti-osteopontin (Biotechne, AF1433).

### Flow cytometry analysis and FACS isolation

Organoids were dissociated in collagenase IV (Worthington, LS004212) at 37°C for 40 min, washed in PBS and digested using a tumor dissociation kit (Miltenyi Biotec, 130-095-929) following the manufacturer’s protocol at 37°C for 30 min. Samples were stained in Zombie Aqua (BioLegend, 423101) at RT for 20 min, after wash and filtration through FACS tubes in PBS. Surface marker staining used the following antibodies: Apoptracker^TM^ Green (BioLegend, 427402), anti-CD68-PE (BioLegend, 333808), anti-CD14-PerCP/Cy5.5 (BioLegend, 324622), anti-CD80-PE/Cy7 (BioLegend, 375408), anti-EpCAM-APC (BioLegend, 324208), anti-CD206-Alexa Fluor 700 (BioLegend, 321132), anti-CD40-APC/Cy7 (BioLegend, 334323), anti-CD11b-BV421 (BioLegend, 301324), anti-CD86-BV605 (BioLegend, 374213), anti-HLA-DR-BV650 (BioLegend, 307650), anti-CD163-BV786 (BioLegend, 333632), anti-CD45-BUV395 (BD Horizon, 563792), anti-CD8-FITC (BioLegend, 301060), anti-CD3-PE (BioLegend, 300408), anti-CD45-APC (BioLegend, 304012), anti-CCR7-Alexa Fluor 700 (BioLegend, 353244), anti-CD4-APC/Cy7 (BioLegend, 300518), anti-CD45RA-BV786 (BioLegend, 304140), anti-CD90-Alexa Fluor 700 (BioLegend, 328120), anti-CD47-BV605 (BioLegend, 323120), anti-CD3-PrCP/Cy5.5 (BioLegend, 300328), all diluted at 1:50 in FACS buffer and stained on ice except for Apoptracker^TM^ Green which was stained at 400 nM. After wash in FACS buffer, the organoid cells were sorted and analyzed in a BD FACSAria-Ⅱ SORP machine after sequential gating for viable cells and singlet cells. Macrophages were gated on CD45+CD11b+HLA-DR+CD68+ cells. We used CD163 to distinguish monocytes (CD45+CD11b+HLA-DR+CD14+CD163-) versus TAM (CD45+CD11b+HLA-DR+CD14+CD163+). Representative gating strategy is shown in **Supplementary Fig. S1A**.

### Bead-based phagocytosis assay using imaging flow cytometry

Single suspension PDO cells (1 x 10^5^) were co-cultured with 1 x 10^5^ streptavidin-coated fluorescent magnetic particles yellow (Spherotech, FSVM-8052-2) in a 96 well ultra-low attachment round bottom plate (Corning, 7007). at 37℃ or 4℃ for 2 h. After co-culture, cells were stained with for FACS using anti-CD45 BUV396, anti-CD11b BV421, HLA-DR BV650 and anti-CD163 BV786. Single cell images were acquired in an imaging cytometer BD FACS Discover S8 sorter. After strictly gating on CD45+CD11b+HLA-DR+CD163+ macrophages, macrophages that further phagocytosed FITC positive beads were enumerated in FlowJo 10.

### pHrodo-based phagocytosis assay

Intact ALI organoids were plated at 50-100 organoids per well in a 96-well flat bottom plate (Corning, 353072) and allowed to adhere overnight. Organoids were incubated with 100 μg/ml pHrodo^TM^ Red E.coli BioParticles^TM^ Conjugate for Phagocytosis (Invitrogen, P35361) ± 1 μM Cytochalasin D (Invitrogen, PHZ1063). Changes in fluorescence were monitored with an Incucyte Live Cell Analysis System.

### Cell-based phagocytosis assay

Single cell suspension ALI PDO cells were incubated with magnetic beads to isolate TAM (CD11b: Miltenyi Biotec, 130-097-142) and tumor cells (EpCAM, Miltenyi Biotec, 130-061-101) by magnetic cell separation following the manufacturer’s protocol. Organoid-derived TAM and tumor cells were pre-labeled with CellTracker^TM^ Deep Red (Thermo Fisher Scientific, C34565) and Calcein AM (Thermo Fisher Scientific, C3100MP) respectively. 1 x 10^4^ of single cell suspension of CD11b+ organoid TAM and EpCAM+ organoid tumor cells were co-cultured and incubated at 37℃ or 4℃ for 2 h in a 96 well ultra-low attachment round bottom plates (Corning, 7007) with IgG1 isotype control or anti-CD47 (clone B6H12) prior to co-culture. After 2 hours incubation, plates were centrifuged after washing with ice cold FACS buffer. Cells were stained with anti-CD45 BUV396, anti-CD11b BV421, anti-HLA-DR BV650 and anti-CD163 BV786, as described above and analyzed by FACS. The fraction of CellTracker^TM^ Deep Red+CD45+CD11b+HLA-DR+CD163+ cells that was additionally positive for Calcein AM was calculated as macrophages that had phagocytosed the Calcein AM(+) tumor cells.

### Cytokine quantification from organoid culture supernatants

Conditioned medium from PDO were collected at day 8 of culture, reflecting 3 days after media change immediately frozen at -80°C. On the day of assay, samples were thawed at RT. Luminex bead-based assays (MILLIPLEX HCYTA-60K-PX48 and MILLIPLEX HCD8MAG15K17PMX, Millipore) were used to quantitate cytokines, and the assay was performed following to manufacturer’s protocol at the Stanford Human Immune Monitoring Center. Secreted SPP1 (osteopontin) was quantified by sandwich ELISA. A commercially available anti-human SPP1 monoclonal (R&D Systems, mAb1433) was used as the capturing antibody and was coated in PBS at 4 μg/ml onto 96-well ELISA plates for 2 hours after blocking with 1% BSA in PBS. Purified recombinant human SPP1 (R&D systems, 1433-OP) was used as standards to construct calibration curves. Culture supernatant from organoids (1:30 and 1:300 dilutions) and standards were diluted with 1% BSA in PBS and incubated in the wells for 2 hours. After washing with 0.05% Tween-20 in PBS, the samples were incubated with biotinylated anti-human SPP1 detection antibody (500 ng/ml, R&D Systems, BAF1433) in PBS with 1% BSA for 1 hour followed by washing and incubation with HRP-streptavidin (200 ng/ml) in PBS with 1% BSA for 1 hour. After washing, tetramethylbenzidine substrate was incubated for 10 minutes followed by the addition of Stop solution (Alpha Diagnostic International) and measurement of absorbance at 450 nm. The concentrations of osteopontin in samples were calculated from the calibration curves of the purified osteopontin standards.

### Cytokine quantitation from human patient plasma

Patient plasma was collected from advanced solid tumors treated with the anti-CD47 blocking antibody magrolimab as part of a phase I clinical trial (NCT02216409)(40). Cytokine (CCL3, CCL4, IFN-gamma, CXCL-9, CXCL-10, and TNF-alpha) concentrations were measured using fit-for-purpose validated Simple Plex^TM^ assays using the Ella^TM^ automated immunoassay platform (Bio-Techne, Minneapolis MN). Briefly, plasma samples, stored at -80°C, were thawed at RT, diluted with an equal volume of assay diluent SD13 (Bio-Techne, Minneapolis, MN) and loaded into wells of the Simple Plex^TM^ cartridge. High and Low control samples were also loaded into each cartridge to monitor assay performance. Cartridges were analyzed on the Ella^TM^ instrument, where analytes are detected using specific capture and fluorescent-labeled detection antibodies in microfluidic channels. The intensity of the fluorescent signal is proportional to the amount of analyte present in the sample. Analyte concentrations are interpolated from a factory-generated standard curve for each cartridge. All data analyzed met pre-specified acceptance criteria. Luminex Discovery Assay (R&D Systems, LXSAH15) was used to quantify SPP1/osteopontin (OPN) from human patient serum (N=6) collected at various timepoints from a phase I magrolimab clinical trial (NCT02216409) (40).

### Single cell RNA-seq analysis

ALI tumor organoid cultures were dissociated 8 days after anti-CD47 or IgG1 isotype control treatment as described above. Fresh tumor tissues were dissociated on the same day of receiving samples. Live CD45+ or CD45+CD11b+HLA-DR+CD68+ organoid cells or fresh tumor tissues were purified by FACS into PBS with 10% FBS and subjected to droplet based scRNA-seq with the 10X Genomics Chromium single cell 5’ platform following to the manufacturer’s protocol using the Chromium NextGEM Single Cell 5’ Kit v2 (PN-1000263) and Library Construction Kit (PN-1000190). Gene expression matrices (GEXs) for each sample were generated using CellRanger (v7.2.0) with the hg38 reference genome for alignment, filtering, barcode counting, and UMI counting. The resulting GEXs were loaded into R (v4.3.3) and converted into Seurat objects using the Seurat package (v5.1.0). Low-quality cells were removed for (1) total number of unique genes per cell fewer than 300 and greater than 3,000-5,000, (2) total UMI counts greater than 10,000-20,000 and (3) greater than 10%-15% of reads mapping to mitochondrial genes.

After filtering, cells from all samples were merged. Next, data were normalized, transformed and scaled using SCTransform v2. Principal component analysis (PCA) was performed, and variation (including sample, group, and tissue type) were corrected using Harmony (v.1.2.1). Cells were clustered using the Louvain algorithm, and dimensionality reduction was conducted via uniform manifold approximation and projection for dimension reduction (UMAP). Macrophages and T cells were identified using well-known marker genes (CD68 and *LYZ* for macrophages and *CD3D* for T cells). Subsets of macrophages and T cells were subsequently extracted and subjected to additional clustering for identification of minor cell populations within each subset. Differential expression analysis was performed on macrophages using the FindMarkers in the Seurat R package. Genes with a log_2_ fold-change greater than 1.0 and adjusted p-values (FDR: False Discovery Rate) below 0.05 were considered significant DEGs in organoid TAM treated by anti-CD47 versus IgG1. To construct single-cell trajectories, the integrated Seurat object was converted into a CellDataSet. Trajectory graphs were computed using the learn graph function in the Monocle3 R package (v 1.3.7) and cells were ordered along the trajectory designating C1Q+ TAMs as the root population. Cell-cell interaction was inferred by CellChat (v 2.1.2), which employs a curated ligand-receptor interaction database. We computed interaction for anti-CD47 and IgG1 treated conditions separately and generated individual CellChat objects for each. Two objects were then merged to identify differential interactions between two experimental groups.

### Bulk RNA-seq analysis

Viable CD45+CD11b+HL-DR+CD68+ cells from organoids were purified through FACS isolation into PBS with 10% FBS. After RNA isolation using the PicoPure RNA isolation kit (Applied Biosystems, KIT0204), isolated RNA sample quality and quantity was assessed using the BioAnalyzer RNA Pico Assay. Library construction was performed following the manufacturer’s protocol for SMART-seq v4 Ultra Low Input RNA kit. Libraries were sequenced on an Illumina NovaSeq X Plus. Raw reads were processed using fastp software to remove adapter sequences, low-quality reads, and ploy-N stretches. The cleaned reads were then aligned to hg38 reference genomes using Hisat2 (v2.0.5). Raw gene expression levels were generated with featureCounts(v1.5.0-p3), and FPKM values were calculated based on gene length and raw reads count. Differential expression analysis was performed using DESeq2 (v1.42.1), and genes with a log_2_ fold-change greater than 1.0 and FDR below 0.05 were considered significant.

Gene set scores for Kyoto Encyclopedia of Genes and Genomes (KEGG) pathways were computed using gsva function from the GSVA R package (v1.50.5). For pathway analysis, significant DEGs identified from single cell and bulk RNA sequencing dataset were ranked by their log_2_ fold-change and used as input for a pre-ranked gene set enrichment analysis. Gene ontology (GO) terms (Biological Process, Molecular Function, and Cellular Component) and KEGG pathways were analyzed using gseGO and gseKEGG functions in the clusterProfiler R packages (v 4.10.1). Gene expression heatmap of selected gene signatures were normalized to average expressions of individual genes.

### Targeted DNA sequencing to detect genomic mutation of fresh tumor and organoids

Genomic DNA was extracted from pairs of fresh tumor and day 8 ALI PDO cultures using the DNA Easy Blood & Tissue Kit (Qiagen, 69506). DNA was submitted to the STAMP (Stanford Actionable Mutation Panel for Solid Tumors) assay. The Stanford Actionable Mutation Panel (STAMP) for solid tumors is a targeted next-generation sequencing assay covering a total of 200 genes with potentially clinically actionable mutations and/or frequently mutated in cancers (60). The workflow includes acoustic sonication of tumor genomic DNA, followed by preparation of sequencing libraries and a target enrichment approach to capture genomic regions of interest for sequencing. The enrichment was performed using custom-designed libraries of capture oligonucleotides that target a specific set of genomic regions. Pooled libraries were sequenced on an Illumina sequencing instrument. Pooled Fastq files were demultiplexed, and reads with non-matching barcodes were discarded. Mapping was performed against the human reference genome hg19, and variants are called separately for single-nucleotide variants (SNVs), indels, fusions, and copy number alterations. Variant calling results were converted to a VCF format, and genotyping and QC reports were generated and reviewed by a board-certified Molecular Genetic Pathologist.

### Machine learning model evaluation of activated microglia

Immunofluorescence image of glioblastoma ALI PDO stained with anti-IBA1 (FUJIFILM Wako, 019-19741) and DAPI were analyzed by a deep learning-based classification model that was developed to distinguish activated and resting microglia based on morphological characteristics. The model was based on EfficientNet, a convolutional neural network family optimized for biomedical image analysis (61). The model’s performance was evaluated using precision-recall metrics and AUC-ROC. Images were processed to determine the total number and proportion of activated and resting microglia. Code is available upon request.

### Statistics and reproducibility

All data are representative of at least 3 biological replicates. Non-parametric two-tailed Mann-Whitney tests were used to determine statistical significance for two samples from different patients, and two-tailed Wilcoxon tests were used for two paired samples from the same patients. *P* values are denoted as \**p* =<0.05,\*\**p* =<0.01 and \*\*\**p* =<0.001.

## Data Availability

This study did not generate new unique reagents. Single-cell RNA-seq (GSE292336) data and RNA-seq data (GSE292328) have been deposited at GEO. All original code has been deposited at https://doi.org/10.5281/zenodo.17087682.

## Author’s Contributions

Conceptualization, MN, CJK. Data curation, LH, RN, JPB. Visualization, LH, RN, MN. Investigation, MN, YLI, YLIU, YPY, MP, LPM, LZ, ET, JP, AF, MFE, KY, CR, HH, RP. Resources, YPY, JO, AB, PD, RM, ATG, JL, EN, ML, CKP, MHG, JTL, LLKL, ASC. Methodology: MN, CJK. Writing – original draft: MN, CJK. Writing – review and editing, MN, RM, JB, CJK. Project administration: RM, MMD, MB, LLKL, JO, JB. Supervision, CJK.

## Acknowledgments

We thank members of the Kuo, Majeti and Bassik laboratories and Kouta Niizuma and Masashi Miyauchi for discussions. We also thank Stanford core facilities for FACS (Catherine Carswell-Crumpton, Cheng Pan, Joe Pasillas), Human Histology (Pauline Chu), Human Immune Monitoring Center (Iris Herschmann, Yael Rosenberg-Hasson, Holden Maecker), Bioinformatics Service Center (bioinformatics services and computing resources) and Tissue Bank (sample provision). These studies were supported by the Japan Society for the Promotion of Science Overseas Research Fellowship (MN), a Uehara Memorial Foundation Research Fellowship (MN), and a Stanford University School of Medicine Dean’s Postdoctoral Fellowship (MN, ET) and a BRAF LGG consortium research fund (CKP), Support was also provided from the National Institutes of Health 5T32DK705648 (JP), R01CA251514 (CJK), U54CA261717 (CJK, CKP), U54CA261719 (CJK), U54CA224081 (CJK) and OT2CA278713 (CJK), the Scientific Foundation of the Spanish Association Against Cancer, AECC (CJK), the PROMINENT team supported by the Cancer Grand Challenges partnership, Cancer Research UK CGCATF-2021/100010 (CJK) and the Stanford Ludwig Center for Cancer Stem Cell Research and Medicine (RM, CJK). Lastly, we are grateful to the participating patients, their family members and the nurses, investigators, and study staff who contributed to this study.

