## Supplementary Data Figures S1-7 for "Organoid modeling of tumor-associated macrophages reveals phagocytosis checkpoint blockade-induced conversion to an immunosuppressive SPP1+ phenotype"

### Supplementary Data Fig.1

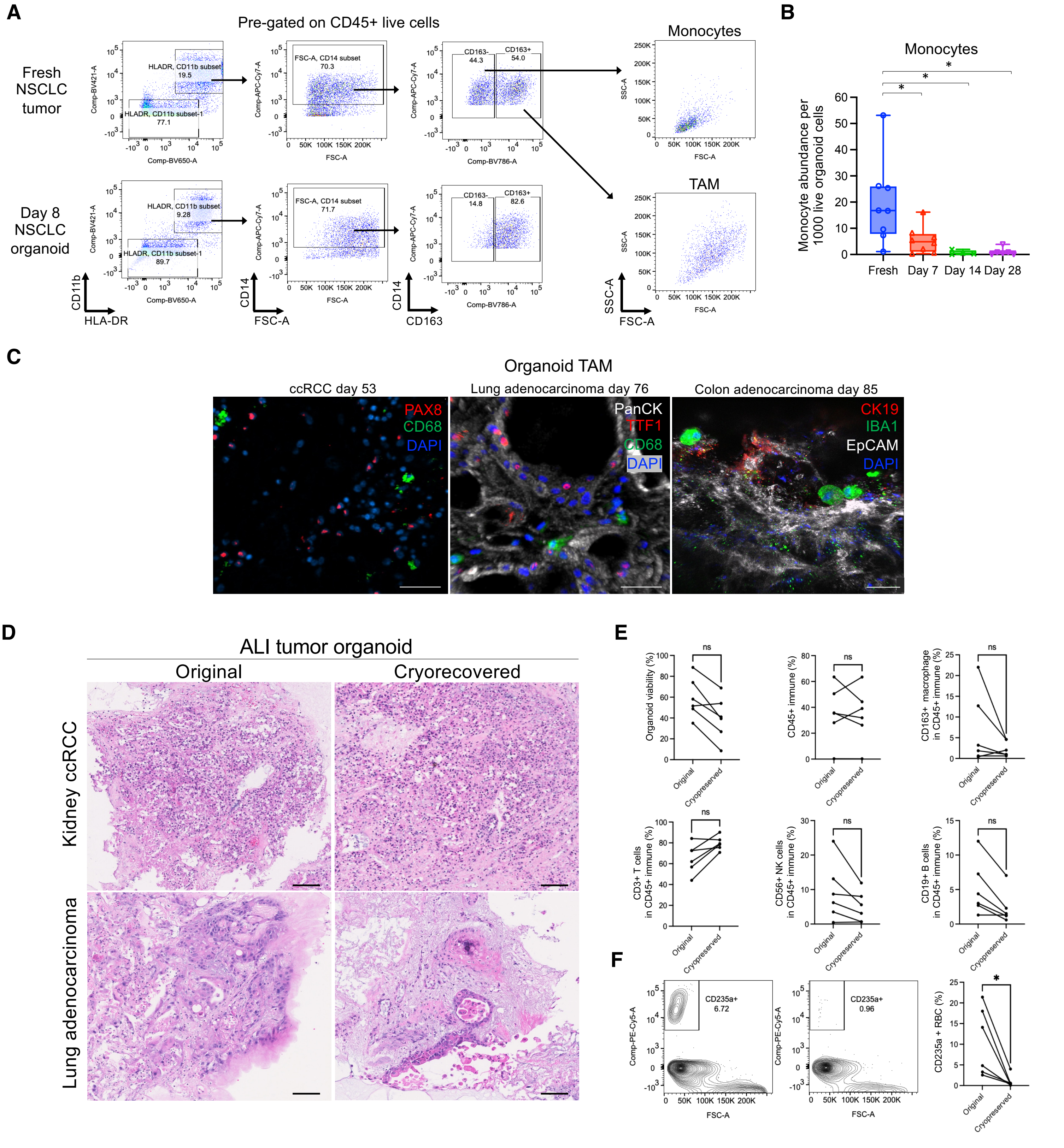


**Supplementary Data Fig. 1. Recapitulation of tumor-associated macrophages (TAM) in ALI tumor organoids.**
**A,** Representative gating strategy for flow cytometry detection of organoid monocytes and TAM. After gating on CD45+ viable immune cells, cells were determined as CD11b+HLA-DR+CD14+CD163- monocytes and CD11b+HLA-DR+CD14+CD163+ TAM. Monocytes and TAM were plotted on forward and side scatter after gating. Fresh tumor and day 8 ALI organoids from the same NSCLC patient are shown. **B,** Box plot showing decreased monocyte abundance over time, flow cytometry from (**A**) (N=8 patients: 5 ccRCC, 2 NSCLC, 1 CRC), **p*=<0.05, Mann-Whitney test. **C,** Representative immunofluorescence image for long term culture ALI tumor organoids showing residual TAM in ALI organoids. TAM were stained with IBA1 (green) or CD68 (green). Tumor epithelium of ccRCC (PAX8: red), NSCLC (TTF1: red and PanCK: white) and CRC (CK19: red and EpCAM: white) were stained with the respective markers. Tumor histologies and culture duration are indicated. All scale bars are 50 µm. **D,** Representative H&E staining of original (left panel) and cryorecovered (right panel) ALI tumor organoids. All scale bars are 100 µm. **E,** Immune subset and viability of original and cryorecovered ALI PDO (N=6 patients: 4 ccRCC, 2 NSCLC). **f,** CD235a positive RBC significantly decreased in cryorecovered ALI PDO (N=6 patients: 4 ccRCC, 2 NSCLC).

Supplementary Data Fig. 2

**A**

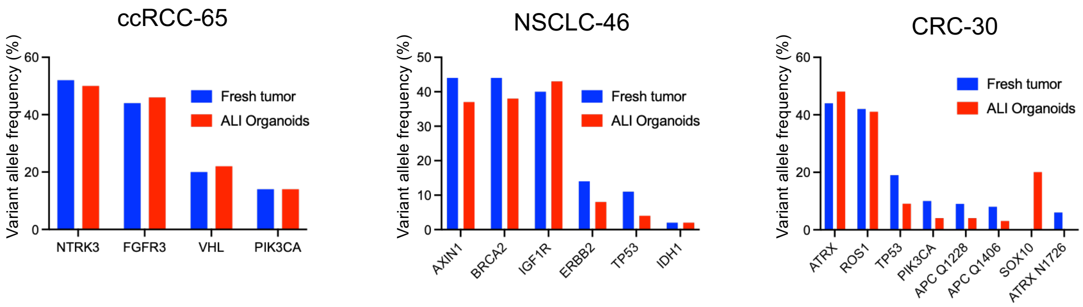


**B**

| Sample ID | Fresh tumor mutation (VAF %) | Organoid mutation (VAF %) |
|---|---|---|
| ccRCC-63 | IKZF3 E277K (46%),VHL E173 (15%),ARID1A R1371Q (13%),GATA3 P135Rfs60 (1%),SUFU P24fs (1%) | IKZF3 E277K (44%),VHL E173 (12%),ARID1A R1371Q (6%) |
| ccRCC-65 | NTRK3 I663V (52%),FGFR3 445+2_445_5del (44%),VHL c.464-1G>A (20%),PIK3A E418K (14%) | NTRK3 I663V (50%),FGFR3 445+2_445_5del (46%),VHL c.464-1G>A (22%),PIK3A E418K (14%) |
| ccRCC-66 | GLI1 R532H (47%),BAP1 c.660-2A>T (22%),VHL E160fs (20%),DNMT3A P904L (2%) | GLI1 R532H (40%),BAP1 c.660-2A>T (15%),VHL E160fs (14%),DNMT3A P904L (4%) |
| ccRCC-67 | SETD2 T305fs (30%),VHL E160fs (26%),SDHB R217C (22%) | SETD2 T305fs (16%),VHL E160fs (16%),SDHB R217C (13%) |
| NSCLC-46 | AXIN1 A526T (44%),BRCA2 A3121V (44%),IGF1R G6R (40%),ERBB2 G778_P780dup (14%),TP53 R273G (11%),IDH1 R132C (2%) | IGF1R G6R (43%),BRCA2 A3121V (38%),AXIN1 A526T (37%),ERBB2 G778_P780dup (8%),TP53 R273G (4%),IDH1 R132C (2%) |
| CRC-30 | ATRX S1046T (44%),ROS1 E1605Q (42%),TP53 E51 (19%),PIK3CA E542K (10%),APC Q1228 (9%),APC Q1406fs (8%),ATRX N1726fs (6%) | ATRX S1046T (48%),ROS1 E1605Q (41%),SOX10 Q364P (20%),TP53 E51 (9%),PIK3CA E542K (4%),APC Q1228 (4%),APCQ1406fs (3%) |
| CRC-31 | LZTR1 A116V (47%),SDHB P254L (44%),PMS2 E710K (9%),TP53 P72fs (7%),APC E1353fs (3%) | LZTR1 A116V (56%),SDHB P254L (50%),TP53 P72fs (21%),APC E1353fs (10%),PMS2 E710K (7%),NOTCH3 G2035fs (1%),HNF1A P291fs (1%) |

**Supplementary Data Fig. 2, Concordance of tumor mutation status in ALI tumor organoids compared with fresh tumor.** Organoid-specific (**A**) and summary table (**B)** of tumor mutation status in representative day 8 ALI tumor organoids compared with fresh tumor in ccRCC, NSCLC and CRC. Variant allele frequencies of respective gene mutations are listed. Targeted DNA sequencing was performed using the Stanford Actionable Mutation Panel for Solid Tumors (STAMP). (N=7 patients: 4 ccRCC, 1 NSCLC, 2 CRC).

Supplementary Data Fig. 3

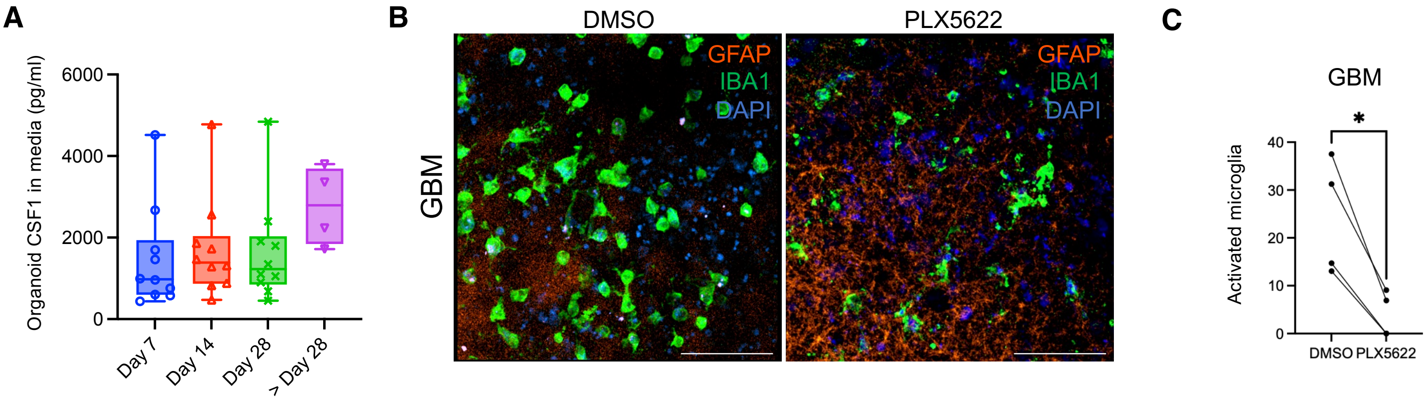


**Supplementary Data Fig. 3. Organoid TAM and microglia are maintained by niche factor CSF-1.**
**A,** CSF1 concentration in ALI PDO media at the indicated time points measured by Luminex (N=14 patients: 6 ccRCC, 3 NSCLC, 5 CRC). Day 29-70 organoids were used as Day>28 ALI PDO. **B,** Representative immunofluorescence staining of day 8 ALI GBM PDO treated with DMSO or CSF-1R inhibitor PLX5622. IBA1 (green), GFAP (orange) and DAPI (blue), scale bar = 50 μm. The GFAP signal could represent either GBM cells or reactive astrocytes. **C,** Quantification of activated microglia with amoeboid histology as a percentage of total IBA1+ microglia from (B) in day 8 GBM organoids, (N=4 patients), *=$p$ <0.05, Wilcoxon test. All scale bars are 50 μm.

Supplementary Data Fig. 4

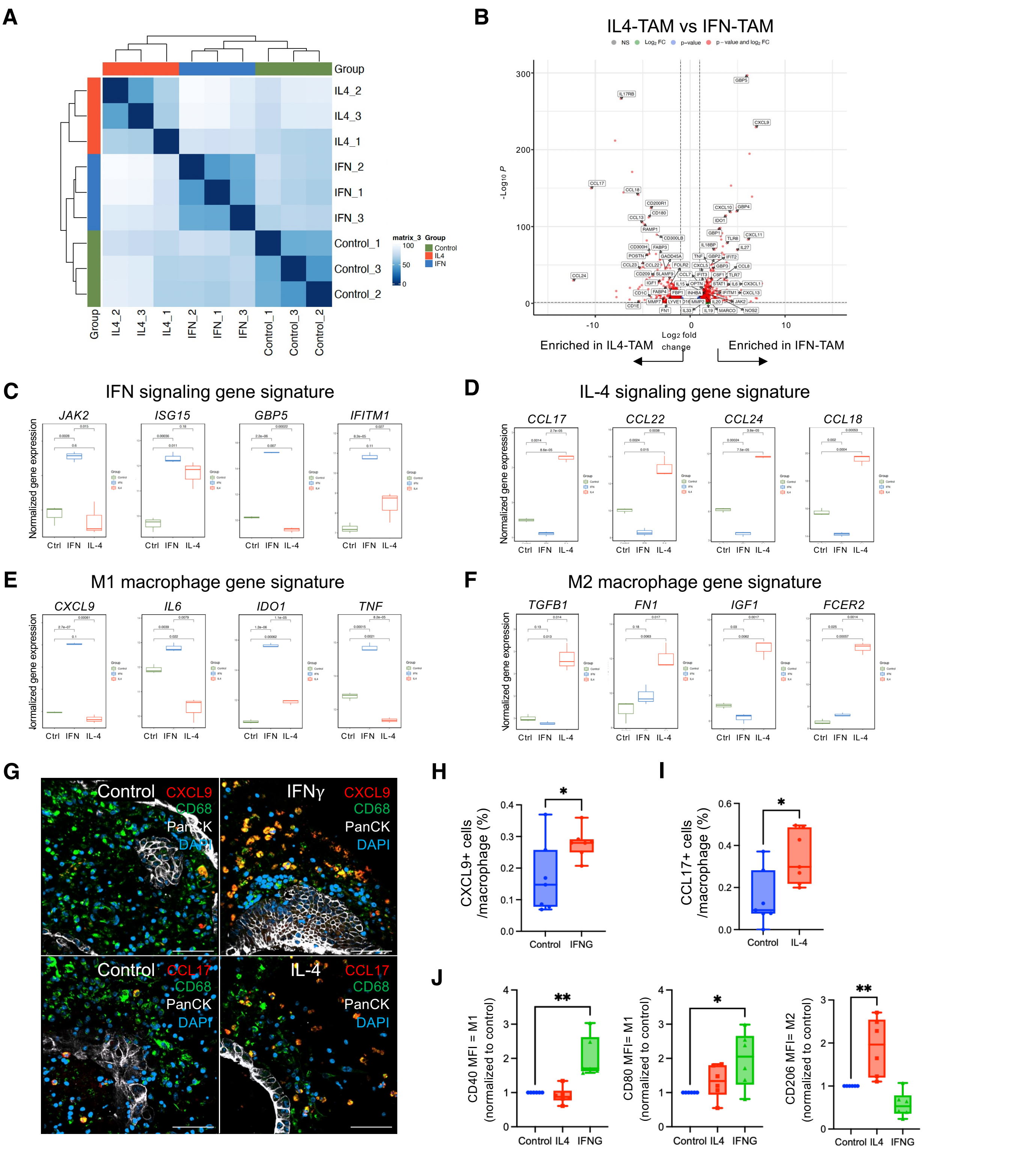


**Supplementary Data Fig. 4. TAM polarization in ALI tumor organoids.**

**A,** Similarity index of bulk RNA-seq transcriptomic data comparing control-, IFNG-, and IL4- treated organoid TAM. n=3 technical replicates from a single ALI ccRCC organoid biological replicate. **B,** Volcano plot showing transcriptomic changes between IFN- and IL4-TAM. $Log_2$ fold-change on the x axis and the $-log_{10}$ adjusted $p$ value on the y-axis. $p$ value of 0.05 and fold-change of 2 are indicated. **C-F,** Box plots of representative gene expression from (**A**) for (**C**) IFN signaling, (**D**) IL-4 signaling, (**E**) M1 and (**F**) M2 macrophages. **G,** Representative immunofluorescence staining showing CCL17 (red), CXCL9 (red), CD68 (green), Pan-CK (white) and DAPI (blue) in ALI CRC organoids, culture day 8, scale bar = 50 μm. **H-I,** Quantification of (**G**) showing CXCL9+ or CCL17+ TAM as a percentage of CD68+ TAM (N=6). **J,** Box plots showing median fluorescence index normalized to control of M1 markers (CD40 and CD80) and M2 marker (CD206) in IFN- and IL-4 treated TAM, measured by flow cytometry-gated CD45+CD11b+HLA-DR+CD163+ cells. N=6, $*p < 0.05$, $**p < 0.01$, Mann-Whitney test.

### Supplementary Data Fig. 5

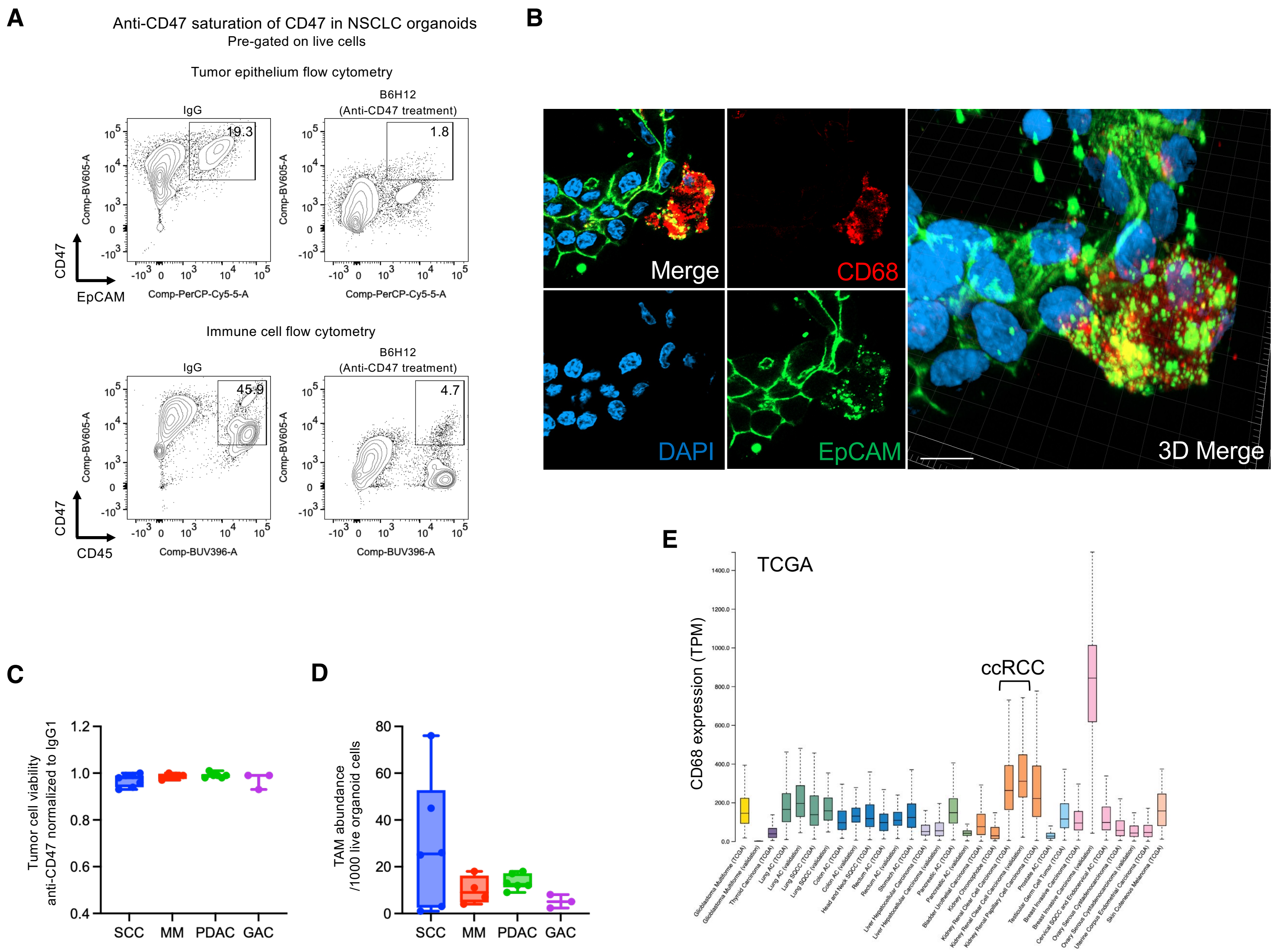


**Supplementary Data Fig. 5. Anti-CD47 treatment induces tumor phagocytosis in ALI organoids.**
**A**, Anti-CD47 saturation of CD47 in NSCLC organoids. Representative flow cytometry plots of ALI tumor organoids (NSCLC) anti-CD47-stained with anti-CD47 detection antibody (anti-CD47-BV605, BioLegend, 323120) after treatment with anti-CD47 blocking antibody (B6H12) or IgG1 treatment, day 8. Panels show CD47 detection with anti-CD47-BV605 in EpCAM+ tumor cells (upper panels) and in CD45+ immune cells (lower panels). The B6H12 anti-CD47 blocking antibody pre-treatment saturates cell surface CD47 and thus strongly blocks CD47 signal from the anti-CD47-BV605 detection antibody (right panels). The CD47 signal in epithelium and CD45+ hematopoietic cells is further consistent with the known ubiquitous expression of CD47. **B,** Representative immunofluorescence of ALI organoid (CRC) showing TAM (CD68, red) phagocytosis of tumor cells (EpCAM, green), scale bar = 10 μm. **C,** Box plots of fractional tumor cell viability in anti-CD47-treated day 8 ALI organoids determined by flow cytometry and normalized to IgG1 control, N=20 patients; 7 cutaneous squamous cell carcinoma (cSCC), 4 malignant melanoma (MM), 6 pancreatic adenocarcinoma (PDAC), 3 gastric adenocarcinoma (GAC). **D,** CD45+CD11b+HLA-DR+CD163+ macrophage abundance per 1000 live organoid cells from (**C**) quantified by flow cytometry (N=20 patients). **E,** *CD68* gene expression values across tumor histologies calculated in TCGA data sets through the Human Protein Atlas website.

Supplementary Data Fig. 6

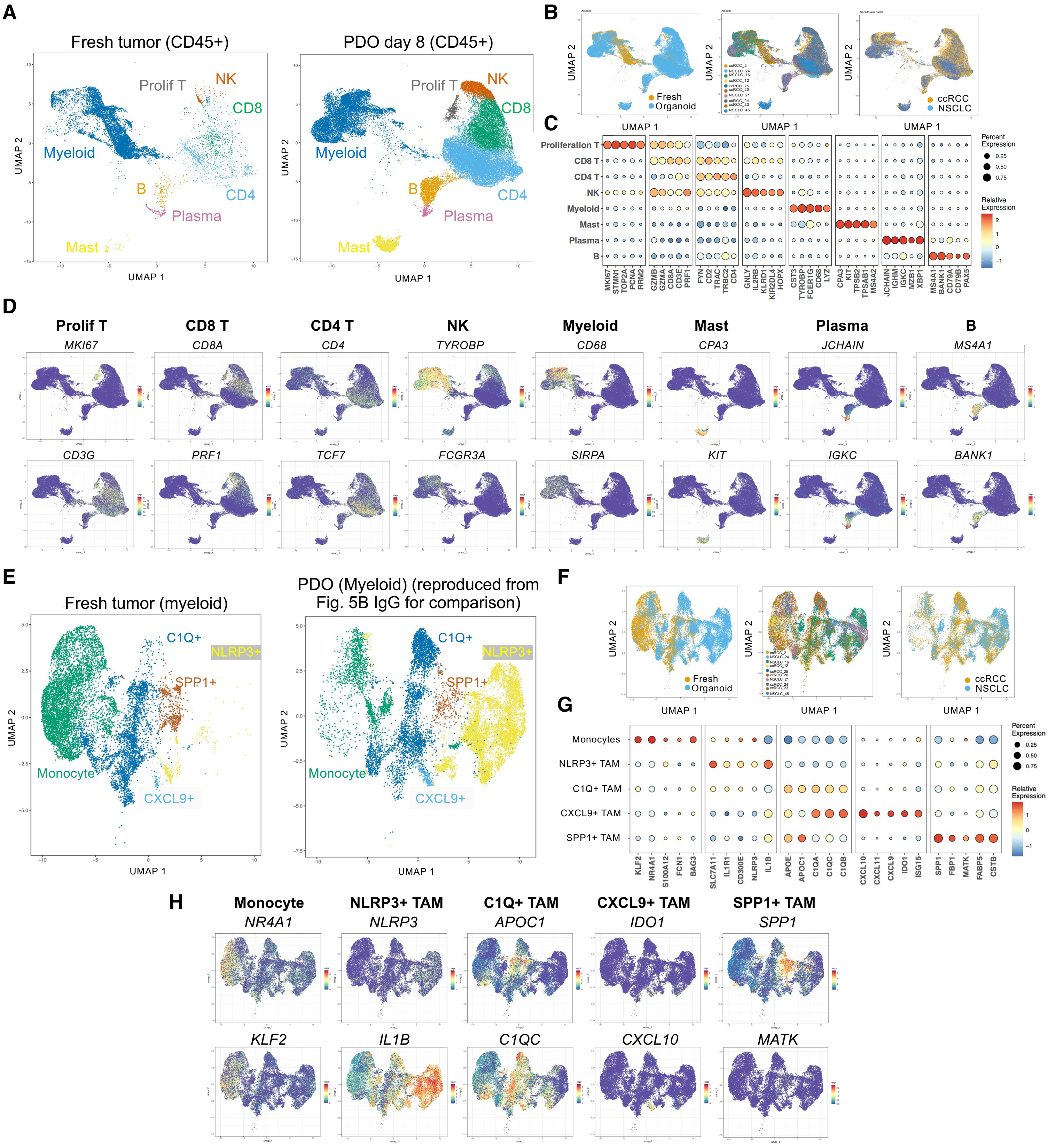


**Supplementary Data Fig. 6. Single cell RNA-seq of immune cells from fresh tumor and ALI tumor organoids.** **A,** UMAP showing major immune subsets from CD45+ FACS-sorted fresh tumor (N=5 ccRCC, N=2 NSCLC) and PDO (IgG condition from Fig. 5A, N=6 ccRCC and N=4 NSCLC). All samples were matched fresh tumor/PDO pairs except for N=1 ccRCC and N=2 NSCLC. **B,** UMAP single cell gene expression analysis from (**A**) in major immune subsets comparing fresh tumor and organoid, by sample ID and tumor type. **C,** Bubble heatmap of expression patterns of selected genes used for cell type identification from (**A**) across indicated major immune subsets. **D,** UMAP feature plots of marker gene expression from (**A**) in major immune subsets. Data is identical to single cell RNA-seq data from Figure 5A (N=10 patients; 6 RCC and 4 NSCLC). **E,** UMAP showing the myeloid (TAM/monocyte, *CD68*+ *LYZ*+) subset from CD45+ FACS-sorted fresh tumor (N=5 ccRCC, N=2 NSCLC) and PDO (IgG condition from Fig. 5B, N=6 ccRCC and N=4 NSCLC). All samples were matched fresh tumor/PDO pairs except for N=1 ccRCC and N=2 NSCLC. **F,** UMAP showing myeloid (TAM/monocyte) single cell gene expression from (**e**) comparing fresh tumor and organoid, by sample ID and tumor type. **G,** Bubble heatmap of expression patterns of selected genes from (**E**) used for cell type identification across indicated TAM/monocyte subsets. **H,** UMAP feature plots of marker gene expression in TAM/monocyte subsets from (**E**).

Supplementary Data Fig. 7

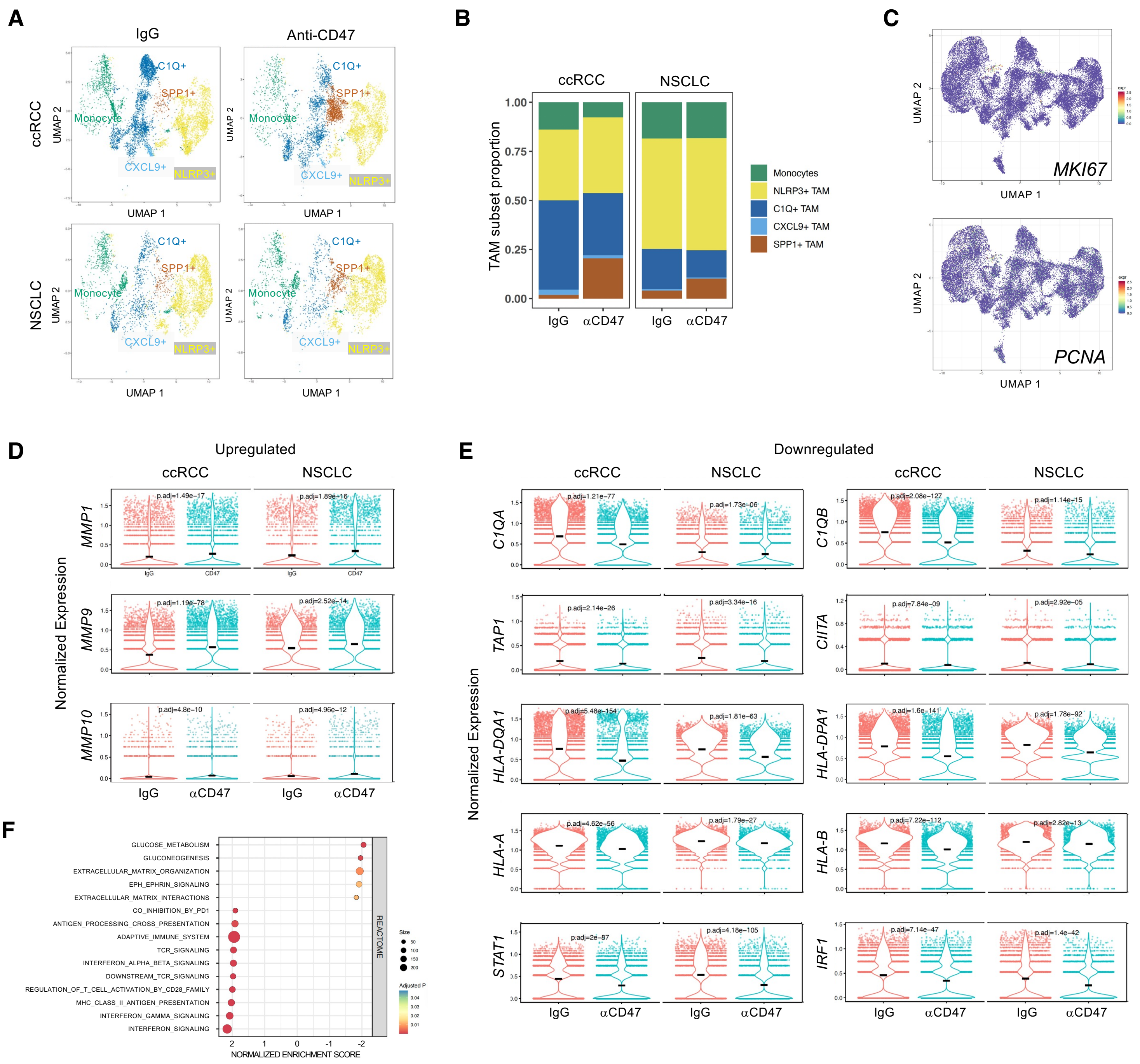


**Supplementary Data Fig. 7. TAM phenotypic changes regulated by anti-CD47 inhibition in ALI tumor organoids.**
**A-B,** UMAP (**A**) and bar chart (**B**) of the TAM/monocyte subsets from Figure 5B scRNA-seq data are shown by tumor type (6 RCC and 4 NSCLC, N=10), depicting PDO TAM subset differences in IgG1- versus anti-CD47-treated day 8 organoids. **C,** UMAP feature plots of proliferation-related gene expression in TAM/monocyte subsets from Figure 5B. **D-E,** Representative scRNA-seq violin plots of gene expression (**D**) upregulated or (**E**) downregulated in anti-CD47 treated ALI tumor organoid TAM. N=10 patients (6 ccRCC and 4 NSCLC) from Figure 5C are shown by tumor types, respectively. **F,** Pathway analysis enriched in IgG1 or anti-CD47 treated ALI tumor organoid TAM from Figure 5C scRNA-seq.
